# Antarctic fish cell cultures show adaptation of organelle morphology and dynamics to extreme cold

**DOI:** 10.64898/2026.04.24.720652

**Authors:** Francesca W van Tartwijk, Anne-Pia M. Marty, Amir Rahmani, Yuetong Jia, Edward N. Ward, Isa Hussain, Lloyd S. Peck, Clemens F. Kaminski, Melody S. Clark

## Abstract

In the Antarctic Southern Ocean, cold-blooded animals have evolved to live at stable temperatures of 0±2 °C. This extremely low temperature affects their biology at every scale, from protein folding to development. However, how animal (sub)cellular organisation and dynamics are adapted to near-0 °C temperatures has not been studied. We therefore established methods to culture and fluorescently label cells from the Antarctic plunderfish *Harpagifer antarcticus* and a temperate comparator species, the shanny *Lipophrys pholis*. By imaging these cultures live at physiological temperatures, we found that subcellular organisation is broadly conserved in *H. antarcticus*, featuring known membranous organelles and biomolecular condensates that remain dynamic, with mitochondria in *H. antarcticus* and *L. pholis* moving at similar speeds. However, we also identified differences in organelle properties between *H. antarcticus* and *L. pholis*, including lysosomal enlargement and mitochondrial morphology changes. These differences may be functionally linked to protein misfolding and slow embryonic development in Antarctic species.

## 1 Introduction

Temperature affects the behaviour of living systems, influencing key cellular processes such as enzyme kinetics, membrane fluidity, and macromolecular assembly ^1, 2^. Cold-blooded animals have therefore evolutionarily adapted their physiology to their habitat temperature, from the whole-body to the cellular and the molecular level. Antarctic marine fauna represent a unique model system to study such adaptation: they have evolved in relative isolation in extremely cold seas for at least 10-15 Myr ^3^, with current temperatures in the Southern Ocean varying between -2 to 2 °C year-round ^4^.

Studies of these Antarctic species have demonstrated that cold-adaptive mechanisms become constrained as water nears its freezing point (<5 °C). At the whole-animal level, polar species’ growth is disproportionately slower than expected from Arrhenius-predicted temperature-dependent slowing of reactions ^5^. At the molecular level, protein folding and proteostatic mechanisms are imperfectly adapted ^5, 6^, as evidenced by observations from the RNA to the protein degradation level ^7–16^. These proteostatic challenges may limit cell proliferation rates and hence may explain the slow growth of polar animals ^5, 6^. However, testing this hypothesis requires model systems of cold-adapted species that permit investigations at the subcellular level near 0 °C, and such systems are currently lacking.

Cold-adapted proteostasis and subcellular organisation likely interact in two ways. Firstly, constitutively high levels of protein misfolding could profoundly affect cytoplasmic organisation, requiring adaptation of the organelles responsible for protein synthesis-related processes such as energy generation (mitochondria) and protein aggregate clearance (the endolysosomal and autophagic systems). Secondly, protein synthesis is spatiotemporally regulated through biomolecular condensation, a temperature-sensitive form of subcellular compartmentation, and it is unknown to what extent this can be maintained in cold-adapted cells ^2^. Therefore, Antarctic animal cell culture methods are needed to enable multi-scale integration of theories on cold adaptation.

Here, we present a method for culturing cells from the Antarctic spiny plunderfish *Harpagifer antarcticus*, as well as from an ecologically similar temperate comparator, the shanny *Lipophrys pholis.* We combine this culture method with live-cell fluorescence microscopy performed near 0 °C ^17^, by testing and adapting fluorescence staining protocols developed for mammalian cell culture to work for *H. antarcticus* and *L. pholis*. We report some of the first insights gained using this model system, on the adaptation of organelle structure and dynamics to extreme cold.

These methods open up a range of novel research directions in cold biology, with significance for cryopreservation and the study of enzymes. Furthermore, as Antarctic marine animals cannot survive long-term at temperatures much above current summer maxima ^18, 19^, this work paves the way towards understanding cellular limitations on the adaptability to global warming of this highly threatened Antarctic ecosystem.

## 2 Results

### 2.1 *H. antarcticus* explants show robust outgrowth of cell monolayers

To establish primary cell cultures from *H. antarcticus*, an explant approach was adopted. Compared with primary cell culture methods using enzymatic dissociation of cells, explantation circumvents challenges with low enzyme activity at low temperature and preserves more native cell-cell interactions and tissue architecture. Small pieces of tissue (∼2 mm x 2 mm) were explanted from fish, placed in 8-well glass-bottom dishes and allowed to adhere before medium addition; after one week, the outgrowth of cells was assessed (Figure 1a). The suitability of several adult tissue types for cell culture was evaluated: kidney, liver, heart, muscle, fin, skin, ovary, and gill. Adherent cell cultures could only be obtained from skin, fin, and ovary (Supplementary Figure 1a). Cultures could also be obtained from embryonic tissue (Supplementary Figure 2). As embryo availability was limited and ovarian tissue could only be obtained from female fish, skin-and fin-derived cultures were primarily used for the method development and experiments reported here.

**Figure 1.**
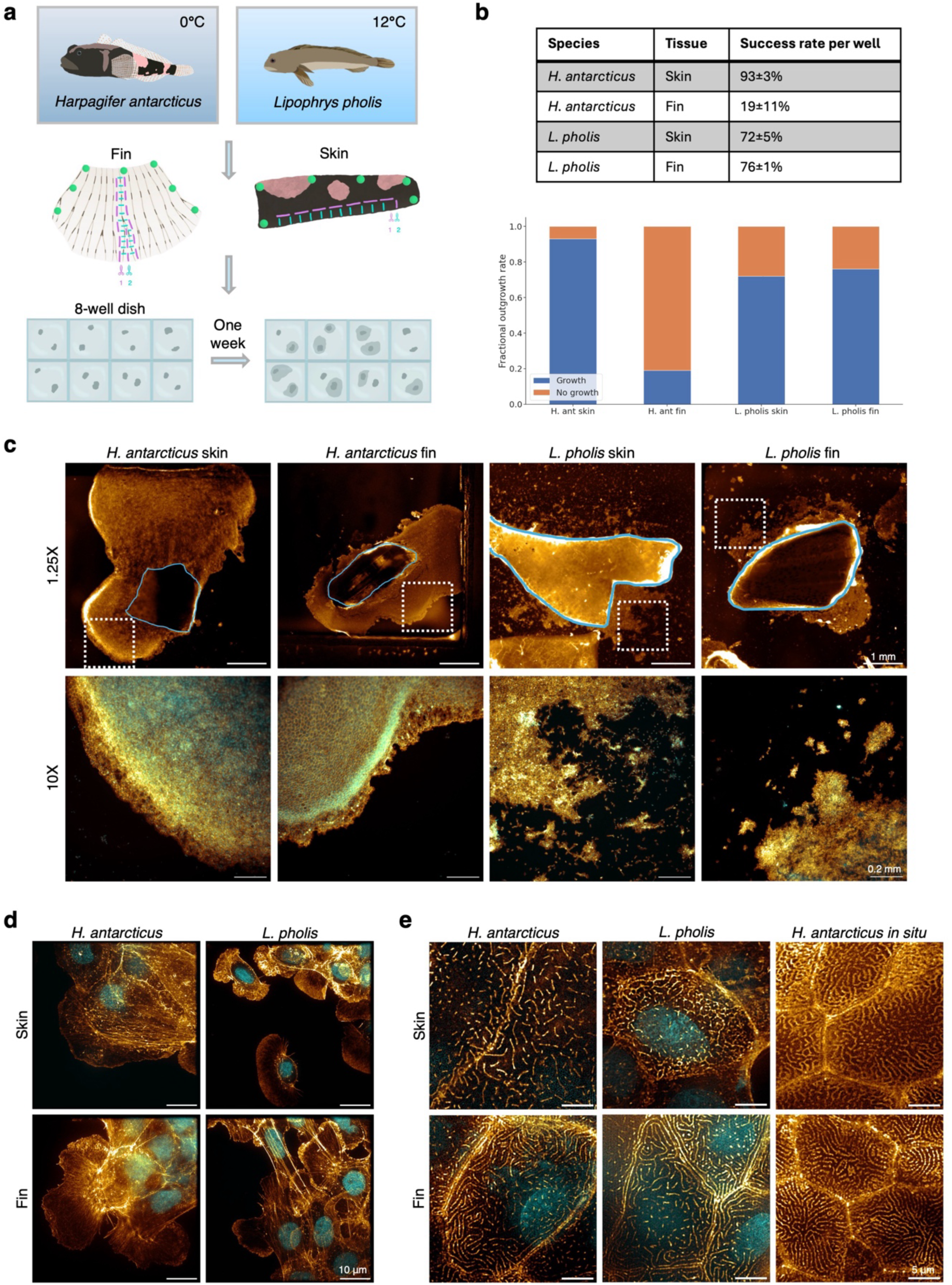
Explantation of skin and fin tissue results in the outgrowth of adherent cells for both *H. antarcticus* and *L. pholis*. a) Schematic overview of explant procedure. Fin and fish tissue were immediately dissected from fish killed according to local regulations. Tissues were immobilised using pins (green) on Sylgard dishes, cut into strips (magenta cut lines), then into rectangles (cyan cuts). Two (skin) or three (fin) pieces of tissue were placed in each well of a culture dish pre-coated with fish-derived gelatin. b) The fraction of wells exhibiting successful cell outgrowth for both species and tissue types (*n*=111 wells (skin) and *n*=72 (fin) for *H. antarcticus*, *n*=112 wells (skin) and *n*=56 (fin) for *L. pholis*, from *N*=3 repeats for all). c) Explant cultures show robust collective migration at one week in culture, with stronger maintenance of epithelial morphology in *H. antarcticus*. Top images: boundaries of explanted tissue are indicated by solid blue line, dashed white squares indicate zoomed-in regions (bottom images). (Widefield fluorescence images) d) At the leading edge, cells have large lamellipodia (images recorded with structured illumination microscopy, SIM). e) Follower cells have an epithelial morphology and display apical microridges. However, microridges are longer *in situ* in *H. antarcticus* fixed native tissue. (SIM) c-e) Samples were fixed, stained with SYTO11 (cyan) and Phalloidin-Atto647N (gold), and imaged at room temperature.

Culture methods were optimised for *H. antarcticus* and their suitability was then tested for *L. pholis*. For the substrate, coating cell culture chambers with commercially available fish-derived gelatin promoted cellular outgrowth (Supplementary Figure 1b), but no other coatings tested did so (see Methods, Fish cell culture). Gelatin was therefore used as a substrate for all cultures in this work. For the culture medium, L-15 supplemented with foetal bovine serum (FBS) and antimicrobials was chosen, as this is common for fish explant cultures ^20^. Further modifications to this medium composition were tested but not incorporated: increasing medium osmolarity through addition of NaCl did not alter cell viability, which was very high at both osmolarities (Supplementary Figure 1c), and substitution of FBS with salmon serum resulted in loss of explant adherence and cell outgrowth (data not shown). Cultures were maintained at 2 °C, to maximise growth rates without causing cell stress: a temperature of 2 °C is within *H. antarcticus*’ natural habitat temperature range ^21^, and has been associated with faster growth (development) of some Antarctic species compared with 0 °C ^22^. With this optimised protocol, using two pieces of tissue per well for skin and three pieces of tissue per well for fin, the fraction of wells with cell outgrowth was very high for skin, and moderate for fin (Figure 1b). For *L. pholis*, following the same protocol resulted in robust outgrowth (Figure 1b), facilitating comparative studies. *L. pholis* cultures were maintained at temperatures to which animals were acclimated prior to capture, which were 6 °C (winter) or 10 °C (spring-autumn).

Following culture method optimisation, cell morphologies were characterised. For both species, fin and skin cells displayed robust outgrowth from the explant within one week, but the nature of their migration was different, as visualised by fixation and staining of filamentous actin (F-actin) and nuclei (Figure 1c). For *H. antarcticus*, fin and skin cells migrated largely collectively: cells formed a sheet extending from the explanted tissue, with almost no single cells occurring. This mode of outgrowth was specific to well-adhered epithelial tissue, as single cells were obtained from ovarian tissue (Supplementary Figure 3). In contrast, *L. pholis* fin and skin cultures showed more jagged outgrowth borders and a greater number of single cells or small clusters of cells. For fin and skin explants from both species (and embryo explants from *H. antarcticus*, Supplementary Figure 2a), cells at the leading edge of the outgrowth were phenotypically distinct from follower cells further back in the sheet. Leading edge cells displayed a highly migratory phenotype, with large lamellipodia (Figure 1d). Similarly, single cells from *L. pholis* showed a migratory keratocyte morphology, also with large lamellipodia. In contrast, follower cells further back in the cell sheet displayed an epithelial morphology, with ‘pebble-like’ cell shapes and (partial) formation of apical microridges (Figure 1e), which are commonly observed for fish and other animal epithelia ^23^. For both species, this explant-derived migration likely reflects a wound healing response, with collective migration of an epithelial sheet that involves a partial epithelial to mesenchymal phenotypic transition, particularly for leading edge cells ^24^. This is supported by the observation that microridges were longer in native tissue than in culture (Figure 1e), as partial microridge disassembly occurs during the wound healing response ^23^. Overall, while a within-culture wound healing response occurs for both species, there are mechanistic differences, with the data suggesting that phenotypic plasticity is lower in *H. antarcticus* than in *L. pholis* (i.e., the epithelial to mesenchymal transition is less complete). This could arise from slower proteomic reprogramming due to low (successful) protein synthesis rates in *H. antarcticus*.

### 2.2 *H. antarcticus* cells contain large perinuclear bodies

Following culture optimisation, we tested various fluorescent labelling methods for the investigation of organelle morphology and distribution in cold-adapted cells. We established labelling protocols using mRNA-based transfection (Supplementary Figure 4) and fluorescent dyes, with the aim to image organelles by live-cell structured illumination microscopy (SIM) using a custom-built cold-imaging platform ^17^. As signal-to-noise ratios following transfection were insufficient for SIM, we visualised total cellular organelle content by performing lipid staining using Nile Red. Nile Red is a solvatochromic lipophilic dye: its excitation and emission spectra peaks are strongly dependent on the polarity of the fluorophore environment, shifting to shorter wavelengths as polarity decreases, e.g., when changing from aqueous to lipid environments ^25, 26^. For example, highly apolar Iipid droplets are excited and emit at shorter wavelengths than organelles bounded by lipid bilayers, allowing their distinction^27^.

In both *H. antarcticus* and *L. pholis* cells, a range of lipid-based organelles were present, which fluoresced when excited at various excitation wavelengths (Figure 2a). As expected, lipid droplets were predominantly visible under 488 nm excitation with detection at 525/30 nm. These were more numerous for *H. antarcticus* but were larger in *L. pholis*, consequently covering a greater total area for *L. pholis* (Figure 2b). At 561 nm with detection at 600/37 nm, multiple cellular organelles became visible, including elongated mitochondria and peripheral tubular endoplasmic reticulum (ER), both indicators of good cell health. These staining patterns were broadly similar to those in a mammalian cell line suitable for SIM (COS-7, African Green Monkey embryonic kidney cells; Supplementary Figure 5a). At 640 nm excitation with detection at 676/26 nm, however, the brightest structures in fish cells were large perinuclear bodies, which were absent in COS-7 cells (Supplementary Figure 5a). These bodies existed in all cells in both species. They were also present in *H. antarcticus* embryonic cells (Supplementary Figure 2b), indicating they do not represent previously described age-associated lysosomal enlargement ^28^. The population of these bodies was highly heterogenous in size and staining pattern, and internal structure was resolvable by SIM for larger bodies in *H. antarcticus* (Figure 2c), indicating a multivesicular nature. The total area of these bodies per cell was greater for *H. antarcticus* than for *L. pholis* (Figure 2d). Therefore, perinuclear multivesicular bodies exist in both fish species, but they are enlarged in *H. antarcticus*.

**Figure 2.**
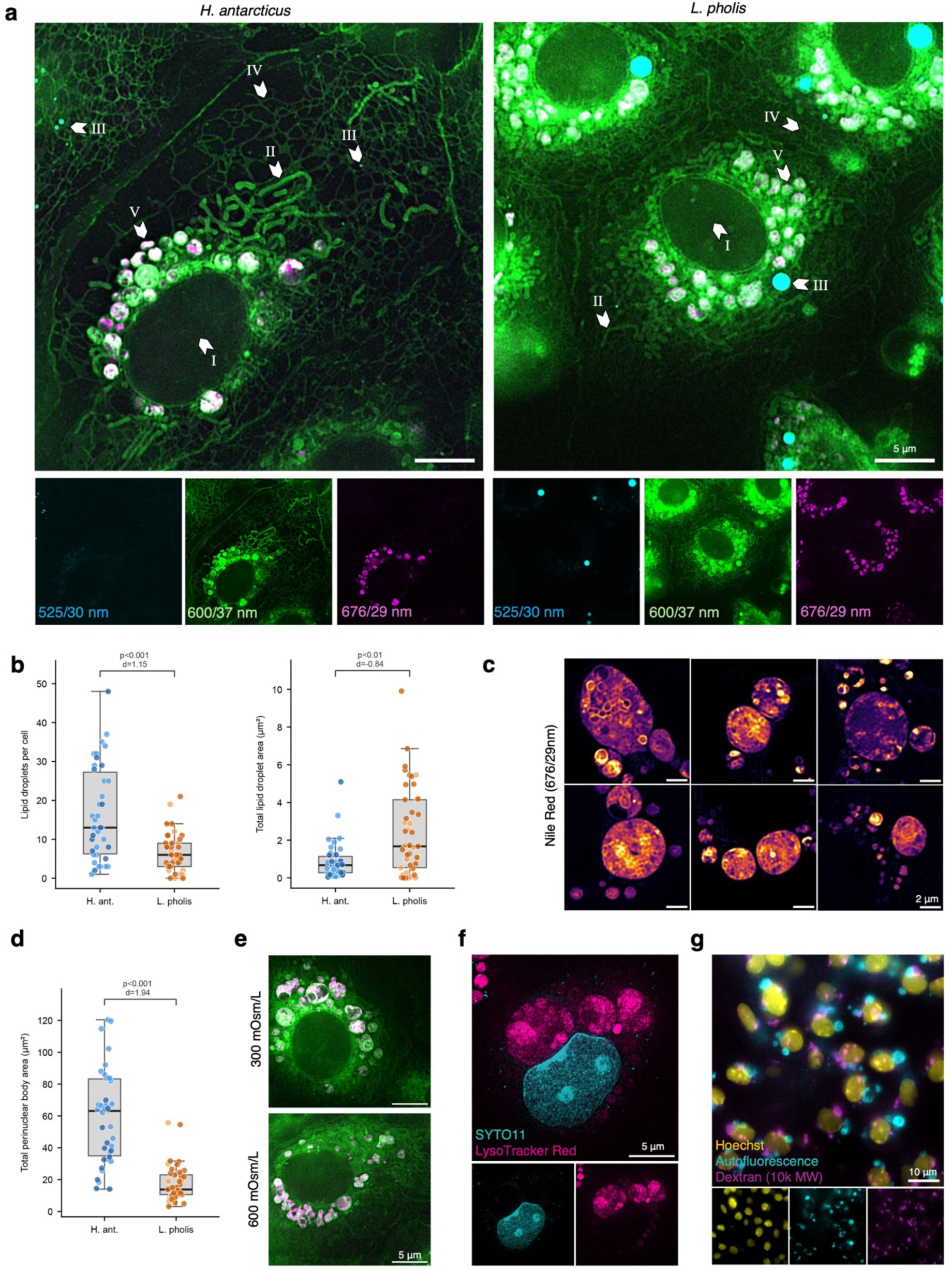
*H. antarcticus* and *L. pholis* contain typical lipid-based organelles as well as large perinuclear bodies. a) Nile Red staining reveals several lipid-based compartments that fluoresce at different emission wavelengths. In the 600/37 nm detection channel (where 600 nm is the central peak of the detection spectrum and 37 nm is the bandwidth), these include I) the nucleus, II) mitochondria, III) lipid droplets, IV) tubular ER, and V) brightly labelled perinuclear bodies. (Example images from fin cell cultures) b) The number of lipid droplets per cell is greater in *H. antarcticus*, but their total area per cell is greater in *L. pholis*. (Fin) c) In the 676/29 nm detection channel, Nile Red mostly stains perinuclear bodies, which are diverse in size and density and show internal structure. (*H. antarcticus* fin) d) The total area per cell of perinuclear bodies is larger for *H. antarcticus* than for *L. pholis*. (Fin) e) *H. antarcticus* skin cells show perinuclear bodies irrespective of medium osmolarity when labelled with Nile Red (600/37 nm emission channel [green] and 676/29 nm emission channel [magenta]). (*H. antarcticus* skin cell cultures derived from the same fish) f) Perinuclear bodies are positive for LysoTracker Red. (*H. antarcticus* skin cells) g) Fluorescently labelled 10k molecular weight Dextran accumulates in perinuclear compartments following overnight incubation. (*H. antarcticus* skin cells) a-e) Imaged live at 2 °C (*H. antarcticus*) or 10 °C (*L. pholis*) using SIM. Median intensity projection of four images. f) Live cell images obtained with SIM performed at 2 °C. g) Fixed-cell images recorded at room temperature on a widefield microscope. b,d) Data points are shaded by replicate.

To investigate the nature of these perinuclear bodies, further experiments were performed using *H. antarcticus* skin cells. To test whether the perinuclear bodies play a role in osmoregulation, cells were grown in media of two different osmolarities. As perinuclear bodies were present under both conditions, their formation is not due to osmotic stress (Figure 2e). The structures were visible when stained with LysoTracker Red (Figure 2f), indicating they have a low internal pH ^29^. They are therefore likely involved in the endocytic and/or autophagic pathways. A Dextran internalisation assay was consequently performed to assess whether these structures contain endocytic cargo. Cells were incubated overnight with 10k molecular weight Alexa647-tagged Dextran, a non-digestible macromolecule, then fixed for imaging. This showed Dextran was internalised into large perinuclear structures (Figure 2g). Additionally, large autofluorescent perinuclear structures were observed in the GFP channel (469/17.5 nm excitation, 525/19.5 nm detection), but these did not overlap with the internalised Dextran signal. Such green autofluorescence is consistent with lysosomal storage bodies ^30^. Therefore, the population of perinuclear bodies likely represents multiple types of endocytic and/autophagic organelles. The largest bodies may represent autolysosomes (formed through fusion of autophagosomes with lysosomes), as these are generally larger than lysosomes ^31^. Irrespectively, as perinuclear bodies are more prominent in *H. antarcticus* than *L. pholis*, these findings indicate cellular ‘digestion’ is less efficient in *H. antarcticus*. This may be related to lower catalytic rates at lower temperatures or to higher loads of misfolded protein aggregates.

### 2.3 *H. antarcticus* mitochondria are abundant, elongated, and motile

To enable further correlation of structure and function in cold-adapted organelle biology, we next sought to selectively label mitochondria: Antarctic fish mitochondria have been studied in isolation and in fixed tissue sections ^32^, enabling functional contextualisation of cell culture-based findings. Cells from both species were labelled using MitoTracker, a dye that requires mitochondrial membrane potential to enter the mitochondrial matrix ^33^, confirming mitochondria remain polarised in culture (Figure 3a).

**Figure 3.**
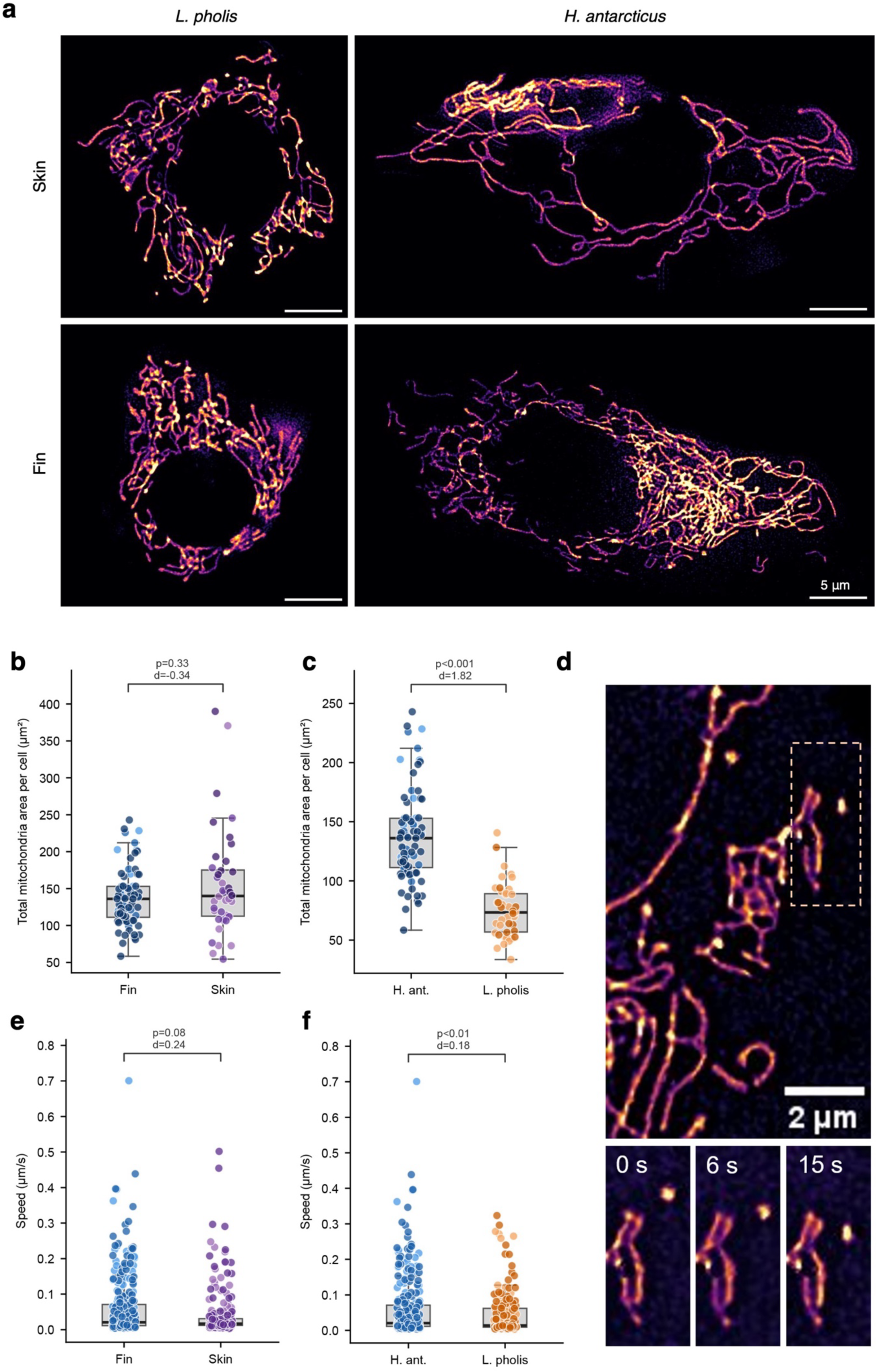
*H. antarcticus* mitochondria display hyperfusion and increased abundance without loss of local motility. a) *H. antarcticus* mitochondria show differences in abundance and morphology compared with *L. pholis*. (For clarity, images have been masked to include only one cell, as for analysis.) b) Total mitochondria area was similar for *H. antarcticus* fin and skin. c) Mitochondria are significantly more abundant in *H. antarcticus* than in *L. pholis* fin. d) Movement of isolated mitochondria is detected by Mitometer: the moving mitochondrion in the zoomed-in region is tracked. e) Mitochondrial speed is similar in *H. antarcticus* fin and skin. f) *H. antarcticus* fin mitochondria show largely similar speed to *L. pholis* fin mitochondria. a,d) Samples stained with MitoTracker Orange. Imaged live at 2 °C using SIM. b,c,e,f) Data points are shaded by replicate.

By observation, mitochondrial abundance and morphology depended on species as well as tissue type (Figure 3a). For *H. antarcticus*, skin cells showed a hyperfused phenotype: mitochondria were highly elongated and/or branched, extending towards the cell periphery. Embryo cultures similarly often showed elongated mitochondria (Supplementary Figure 2c). In *H. antarcticus* fin cells, mitochondria were also elongated but less branched, and very abundant. In contrast, *L. pholis* cells showed a mostly perinuclear distribution of shorter, less abundant mitochondria.

To quantify phenotypic differences, MitoTracker images were segmented for morphological analysis. We compared mitochondrial properties in both tissues for *H. antarcticus* and compared fin samples for interspecies comparison, given the fin cultures’ relatively similar mitochondrial morphology. We used Nellie ^34^ for segmentation, an algorithm which analyses geometric shapes and structures across multiple scales and enhances their contrast before segmentation without relying on image brightness. We also developed our own, brightness-based segmentation pipeline (see Methods, Image analysis). Both analysis methods performed similarly, but the extensive overlap between mitochondria prevented their individual discrimination for subsequent analysis (Supplementary Figure 6). As measures of mitochondrial morphology like eccentricity or form factor rely on accurate segmentation, mitochondrial morphology was not quantified further.

Instead, we quantified total mitochondrial area as a proxy for total mitochondrial mass. Mitochondrial mass thus determined was found to be similar for *H. antarcticus* skin and fin (Figure 3b), and significantly greater for *H. antarcticus* fin compared with *L. pholis* fin (Figure 3c). It was known that highly active red muscle requires higher mitochondrial densities to meet energy demand at low temperature in Antarctic vertebrates: very high mitochondrial densities have previously been observed in Antarctic pectoral adductor muscle fibre ^35, 36^. From our data, it appears that cold adaptation also leads to an upregulation of mitochondrial mass in fin and possibly skin epithelial cells, which have lower respiratory demands. This demonstrates that mitochondrial mass is generally upregulated in cold-adapted cells where cell activity levels are maintained. This may compensate for lower mitochondrial respiratory activity per unit of mitochondrial volume or reflect a general increased demand for ATP to compensate for low-temperature effects.

We next quantified mitochondrial dynamics using the open-source software Mitometer ^37^. For extensively overlapping mitochondria, detection of displacement by Mitometer was challenging; the resulting measurements of speed therefore largely represent the motion of smaller, more isolated mitochondria (Figure 3d). For *H. antarcticus*, mitochondrial speed was similar in skin and fin tissues (Figure 3e). Surprisingly, mitochondrial speed was greater for *H. antarcticus* fin than for *L. pholis* fin (Figure 3f). However, while this effect was statistically significant, the effect size as measured by Cohen’s *d* was very small (*d*<0.2), indicating the effect may not be physiologically significant. Therefore, our data indicate the motility of mitochondria may be highly cold-adapted, being functionally similar in both species.

### 2.4 *H. antarcticus* cells contain nucleic acid-based granules

Finally, we looked to establish strategies to label membraneless compartments in our cell cultures. To label RNA-containing condensates, we used the SYTO Green family of DNA/RNA dyes in live cells. Different SYTO dyes gave different staining patterns in trials using COS-7 cells. Ιn subsequent experiments with fish cells, we used SYTO24 and SYTO12, as these most strongly labelled differently localised granules in COS-7 cells: mitochondrial DNA/RNA-containing granules (Supplementary Figure 4b) and non-mitochondrial non-nuclear RNA granules and the nucleolus (Supplementary Figure 4c) respectively. SYTO24’s mitochondrial staining is expected: the dye has a net positive charge at neutral pH, causing accumulation in mitochondria due to their negative-inside membrane potential.

For both *H. antarcticus* and *L. pholis* skin and fin, SYTO24 strongly labelled intramitochondrial puncta (Figure 4a), as was the case in *H. antarcticus* embryonic cells (Supplementary Figure 2d). These puncta most likely represent mitochondrial nucleoids, mitochondrial RNA processing granules and/or RNA translation hubs, which have been proposed to form through biomolecular condensation and to spatially organise mitochondrial gene expression ^38^. The number of intramitochondrial puncta was lower than typically observed in mammalian cells, in our observations via staining with SYTO24 (Supplementary Figure 4b) or as reported in other studies using other labels ^39, 40^. Some granules localised to mitochondrial tips and branch points, and near sites of fusion, as previously reported for mammalian cells ^40^, as well as near sites of fission (Figure 4b, Supplementary Video 1). In addition, small and dim SYTO24-positive puncta were detectable outside mitochondria, which presumably represent cytoplasmic ribonucleoprotein granules (RNPs) formed by biomolecular condensation (Figure 4c). These puncta displayed directional movement consistent with active transport. These results indicate that even in extremely cold environments, animal cells still rely on DNA/RNA-based biomolecular condensation as a mechanism to organise their internal functions. However, intramitochondrial condensates likely differ from those in mammalian cells in abundance, size distribution, and/or clustering, which could not be fully resolved by SIM.

**Figure 4.**
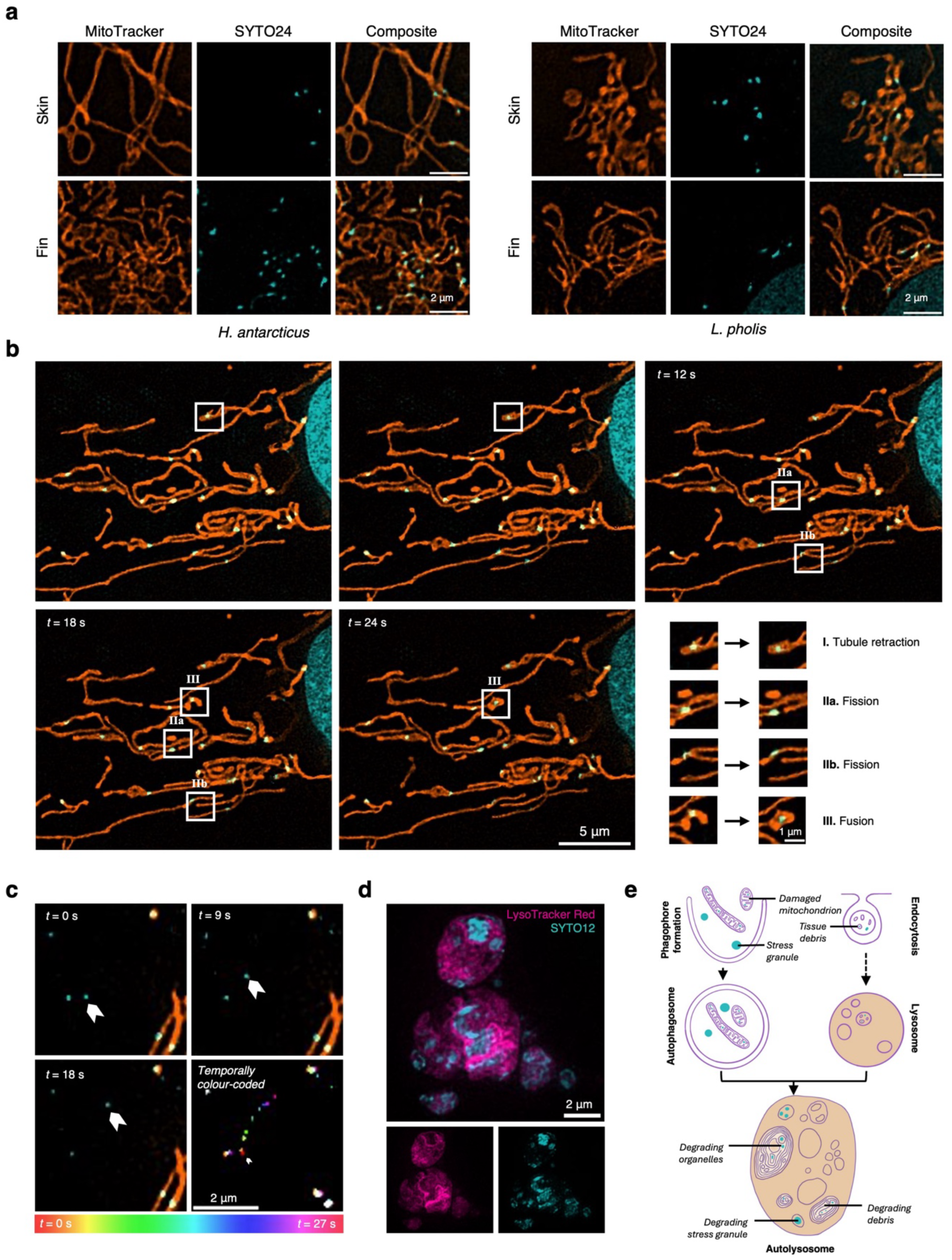
*H. antarcticus* cells contain nucleic acid-based granules. a) Both *H. antarcticus* and *L. pholis* fin and skin cells contain intramitochondrial granules that are labelled by SYTO24. b) Example *H. antarcticus* skin cell in which granules are positioned at sites where mitochondrial remodelling takes place (see zoomed-in detail shown in panels I-III). c) In some cells, SYTO24-positive puncta can be observed within the cytoplasm. The motile punctum shown here (white arrowheads) has an average speed of 0.08 μm/s and a maximum frame-to-frame velocity of 0.16 μm/s. A temporally colour-coded image displaying the complete trajectory is shown. d) SYTO12 stains LysoTracker-positive perinuclear bodies. (*H. antarcticus* skin cell.) e) Nucleic acids could enter autolysosomes through autophagy of stress granules or mitochondria, or through endocytosis of tissue debris. Colour code: membranes in purple, nucleic acids in cyan, acidic internal environments in orange. a-d) Imaged live at 2 °C (*H. antarcticus*) or 10 °C (*L. pholis*) using SIM.

In contrast, when tested in *H. antarcticus*, SYTO12 strongly labelled only the previously identified perinuclear bodies (Figure 4d), not the nucleolus. As the perinuclear bodies are LysoTracker-positive, they are acidic; therefore, SYTO12 accumulation within these structures is not due to the dye’s net positive charge (c.f. SYTO24’s preferential staining of intramitochondrial granules). As for Nile Red and Lysotracker, there was inhomogeneity in SYTO12 staining within single perinuclear bodies: SYTO12 staining was brightest in regions within perinuclear bodies that were LysoTracker-depleted, indicating the SYTO12-positive regions are relatively dense, largely still undigested cargo. This result indicates perinuclear bodies contain nucleic acid-based structures, potentially RNA given SYTO12’s strong nucleolar staining in mammalian cells. This could arise through autophagy of RNA-containing structures like stress granules ^41^ or mitochondria, or through endocytosis of debris from other cells (Figure 4e). The former could indicate challenges in maintaining (the correct phase of) RNA-containing condensates in the cold.

## 3 Discussion

Using our cell culture method, we have presented the first super-resolution images of major cellular organelles in living cells derived from an animal species adapted to extreme cold. Some subcellular features show remarkable levels of conservation compared with temperate species. Our results show that despite the very slow timescales of whole-animal processes such as development, cold-adapted animals remain dynamic at the subcellular level, at least for some processes, with surprisingly high speeds of mitochondrial movement. Furthermore, we show that small nucleic-acid containing granules can exist at low temperature, showing conserved functionality via the process of phase separation.

However, by comparison with a temperate species, we also identify unique subcellular features of cold-adapted cells. We show mitochondrial hyperfusion and (auto)lysosomal enlargement, which may represent functional consequences of challenges in proteostasis ^7–15^. Perinuclear body enlargement may function to degrade clusters of misfolded proteins ^42^, potentially including entrapped RNA if RNA-binding proteins misfold (causing SYTO12-positive staining). Furthermore, mitochondrial hyperfusion may represent a mitigatory response to inefficient cytosolic protein synthesis ^43, 44^ and/or higher levels of reactive oxygen species ^45, 46^ damaging proteins, by increasing mitochondrial ATP production ^43^ and protecting mitochondria from autophagy ^47, 48^. We also demonstrate mitochondrial density is increased in non-muscle cell types. This may compensate for the physical effects of low temperature on molecular motion and catalysis; mitochondrial proliferation may facilitate oxygen diffusion through lipid environments ^49^ and reduce stochasticity affecting intramitochondrial reaction rates at low temperature ^50, 51^. Further research is required to test these and related hypotheses.

We anticipate that our method’s application will lead to further insights in cold biology across biological length scales. Experiments that are now becoming possible include protein folding rate quantification using established methods such as SunTag ^52^ or single molecule translation imaging ^53^, single particle tracking-based characterisation of cytoplasmic biophysical parameters, dynamic localisation of proteins, quantification of cell stress using organelle morphology ^54^, and measurements of cellular division and growth rates. Such experiments will enable us to explore the extent to which cell-autonomous factors as opposed to system-wide properties (e.g., circulatory ability to deliver oxygen ^55^) contribute to whole-animal slow growth rates and low upper temperature limits ^5, 6^. This will inform our understanding of adaptation to extreme cold environments and the vulnerability of Antarctic species to their changing habitat.

## 4 Acknowledgements

This work was supported by a UKRI Cross Research Council Responsive Mode (CRCRM) award [MR/Z505341/1] (MSC and CFK, supporting FWvT and AR), the UK Engineering and Physical Sciences Research Council (EPSRC) [G132372 (APMM), EP/L015889/1 and EP/H018301/1 (CFK)], the Wellcome Trust [3-3249/Z/16/Z and 089703/Z/09/Z] (CFK), the Medical Research Council (MRC) [MR/K015850/1] (CFK), Infinitus China Ltd (CFK), and UKRI-NERC core funding to the British Antarctic Survey (MSC and LSP).

The authors thank the Rothera marine team for collecting *H. antarcticus* and Berwyn Roberts and Laura Grange (School of Ocean Sciences, University of Bangor) for collecting *L. pholis*, and BAS Marine Aquarium Manager Samantha Kentwell for all fish husbandry.

For the purpose of open access, the authors have applied a CC-BY license to any author-accepted manuscript arising from this submission.

Data access statement: all images and code used to generate figures in this work will be made available upon final publication.

## 5 Supplementary figures

**Supplementary Figure 1.**
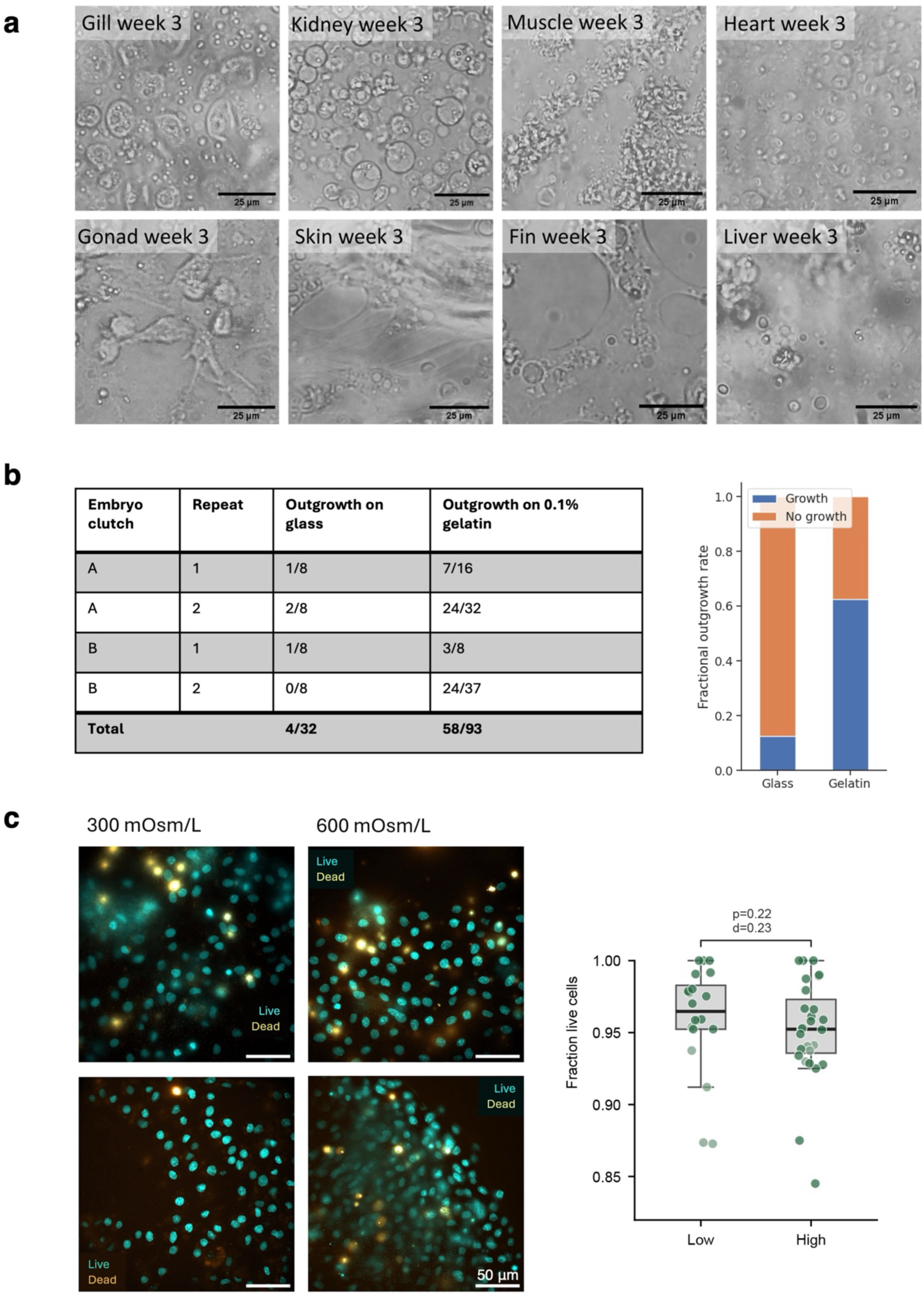
Optimisation of culture protocols for *H. antarcticus* involved trialling different tissue types, medium compositions, and dish coatings. a) Attempts at growing fin, gill, gonad, heart, kidney, liver and muscle from *H. antarcticus* adult tissue types show that only skin, fin, and ovary (which can only be obtained from half the fish dissected) are suitable for 2D cell culture compatible with high-resolution imaging. b) Coating with 0.1% fish-derived gelatin improves outgrowth rates for embryo explant cultures. Cultures were generated from two clutches of embryos in two different experiments. The number of wells in which an explant showed cell outgrowth was quantified. Tabular data are presented as [number of wells with outgrowth]/[total number of wells]. c) Changing medium osmolarity does not alter cell viability. Viability was compared in L-15 at standard osmolarity (300 mOsm/L) and at osmolarity equivalent to that of sea water (600 mOsm/L). Samples were stained with nuclear dyes Hoechst (all cells) and DRAQ7 (dead cells). Data points are shaded by replicate. (Widefield. Imaged live at room temperature immediately after staining.)

**Supplementary Figure 2.**
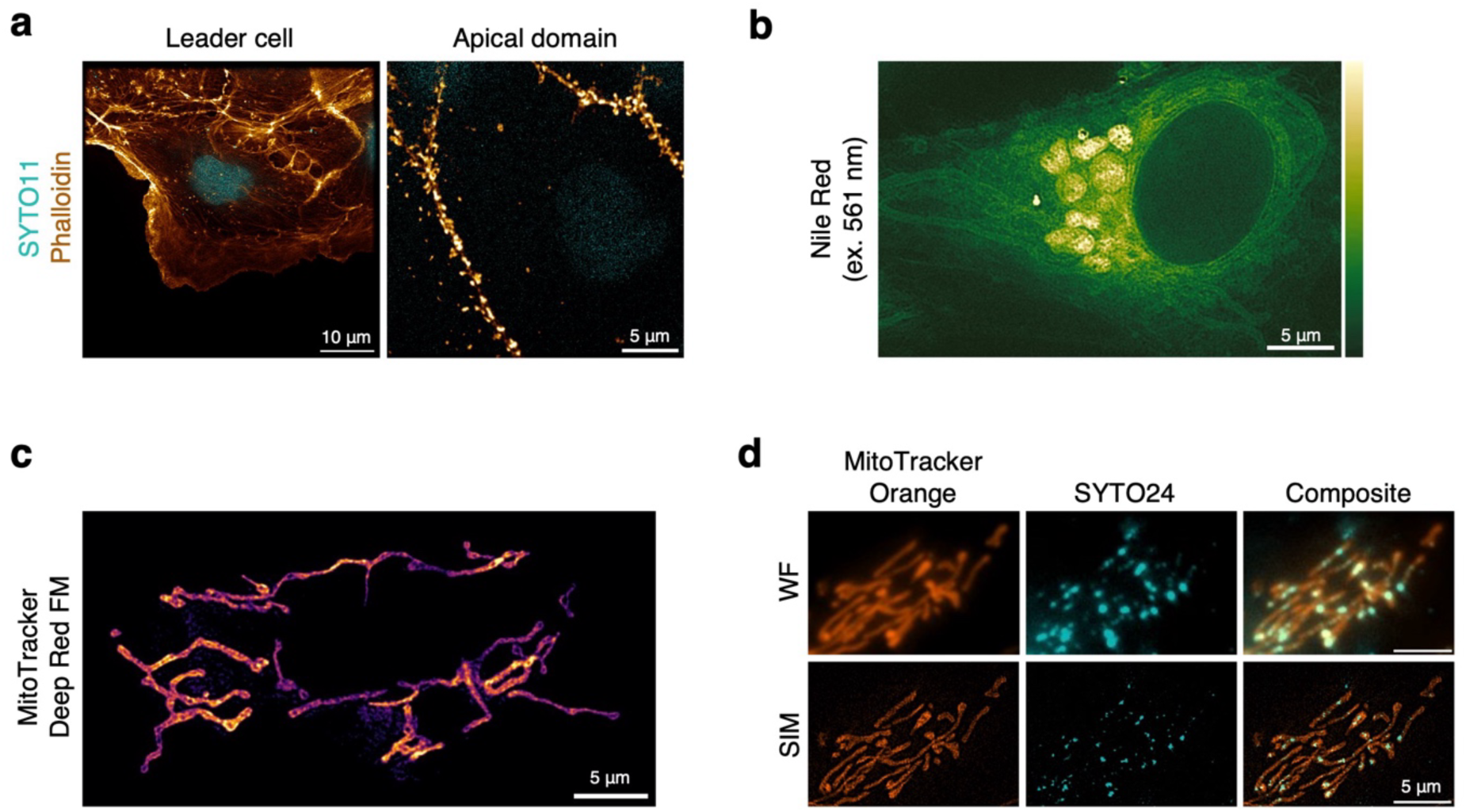
*H. antarcticus* embryo cell cultures show similar subcellular organisation to fin and skin cell cultures from adult tissue. a) *H. antarcticus* embryo cells display outgrowth broadly similar to that of skin and fin cultures. Clear leader cells with large lamellipodia are distinguishable at the outgrowth edge (left). Apical actin patterns are distinguishable in follower cells, though not well-formed microridges (right). b) Embryo cells also contain perinuclear bodies. c) Embryo cells contain branched and elongated mitochondria. d) Embryo cell mitochondria contain nucleic acid puncta. a) Samples were fixed and stained with SYTO11 (cyan) and Phalloidin-Atto647N (gold). Imaged at room temperature using SIM. b-d) Imaged live at 2 °C using SIM. For b, a median projection of four images is shown. For d, both reconstructed widefield and super-resolution images are shown.

**Supplementary Figure 3.**
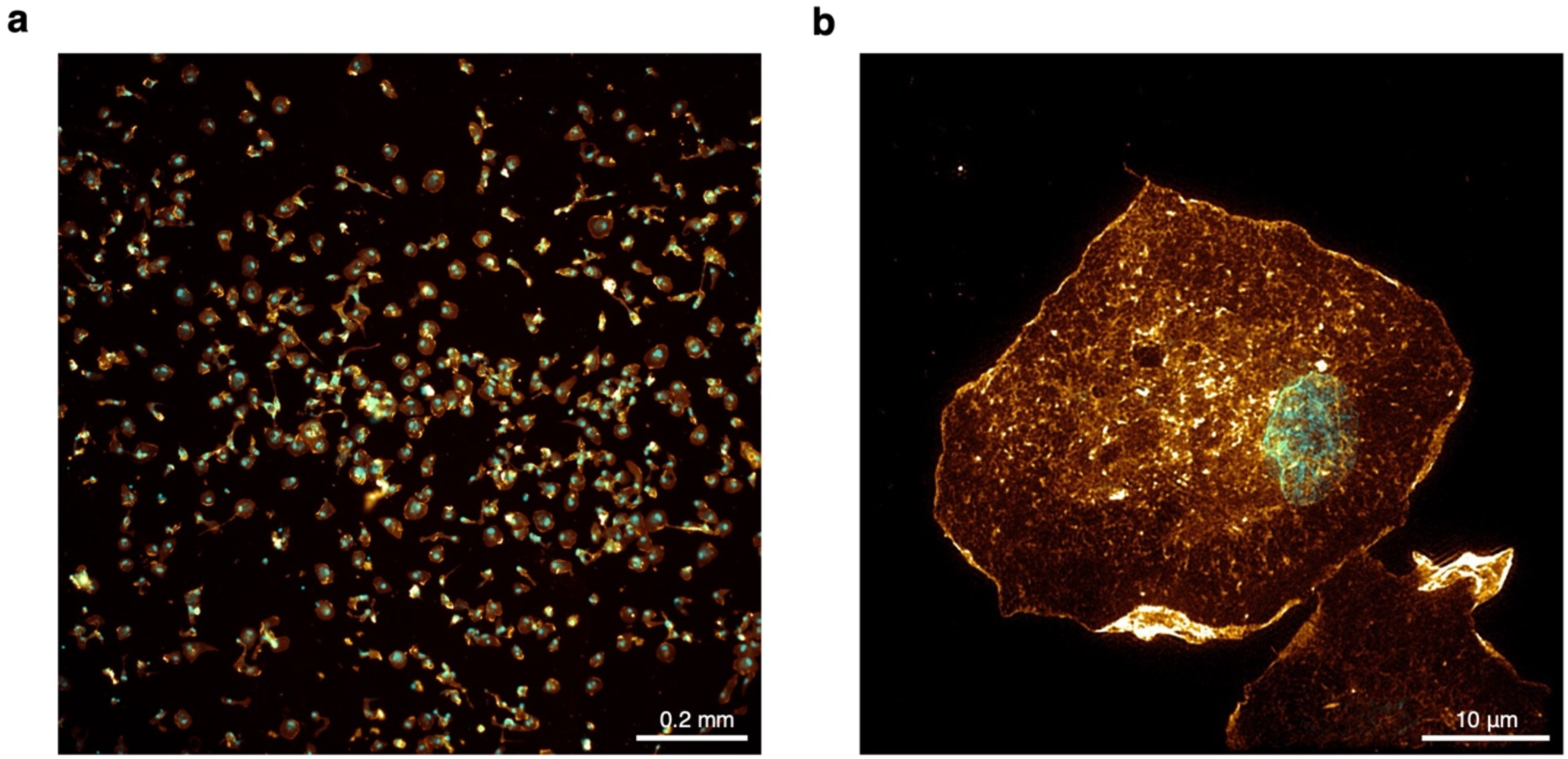
*H. antarcticus* ovarian cultures largely grow as single unpolarised cells. a) Ovarian cells display some clustering but largely grow as single cells, with variable morphology. b) Many ovarian cells show limited polarisation. Samples were fixed and stained with SYTO11 (cyan) and Phalloidin-Atto647N (gold). Imaged at room temperature using widefield (a) or SIM (b).

**Supplementary Figure 4.**
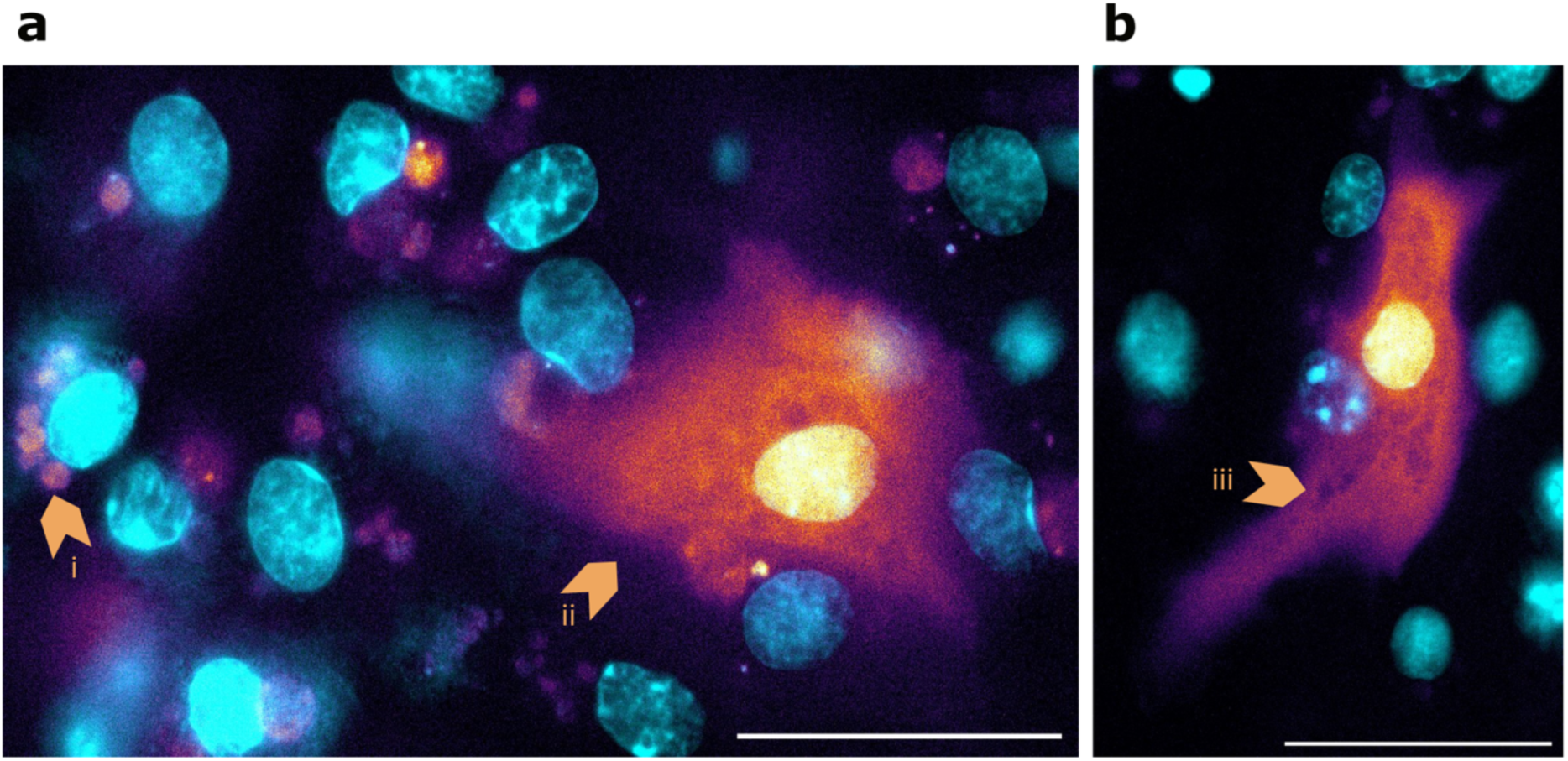
*H. antarcticus* skin cultures can be transfected using mRNA. a) The transfection yield is variable depending on cell confluence. b) In some cases, the labelling density reveals organelles and cytosolic structures. Samples are stained with Hoechst (cyan) and transfected using GFP mRNA (inferno). Arrowheads show (i) autofluorescence in the GFP channel, (ii) a transfected cell and (iii) example of an organelle unlabelled by cytoplasmic GFP. Scale bar is 50µm in both images.

**Supplementary Figure 5.**
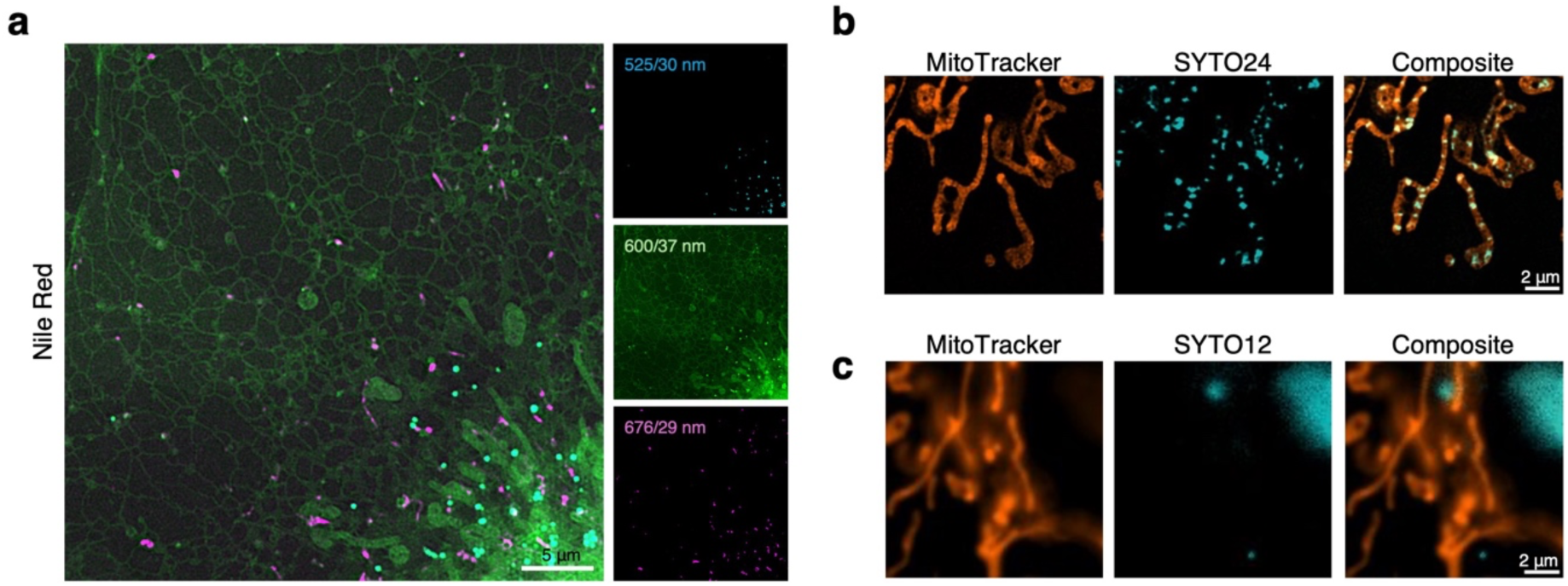
For select dyes, initial staining trials were performed using COS-7 cells. a) Nile Red staining shows lipid droplets at 488 nm excitation with detection at 525/30 nm, a range of organelles including ER and mitochondria at 561 nm excitation with detection at 600/37 nm, and small dynamic vesicles at 640 nm excitation with detection at 676/29 nm. b) SYTO24 strongly labels intramitochondrial nucleic acids. c) SYTO12 labels the nucleolus and what appear to be cytoplasmic RNA granules. a-c) Imaged live at 37 °C with CO2 supply using SIM. For a, a median project of four images is shown. c is a reconstructed widefield image.

**Supplementary Figure 6.**
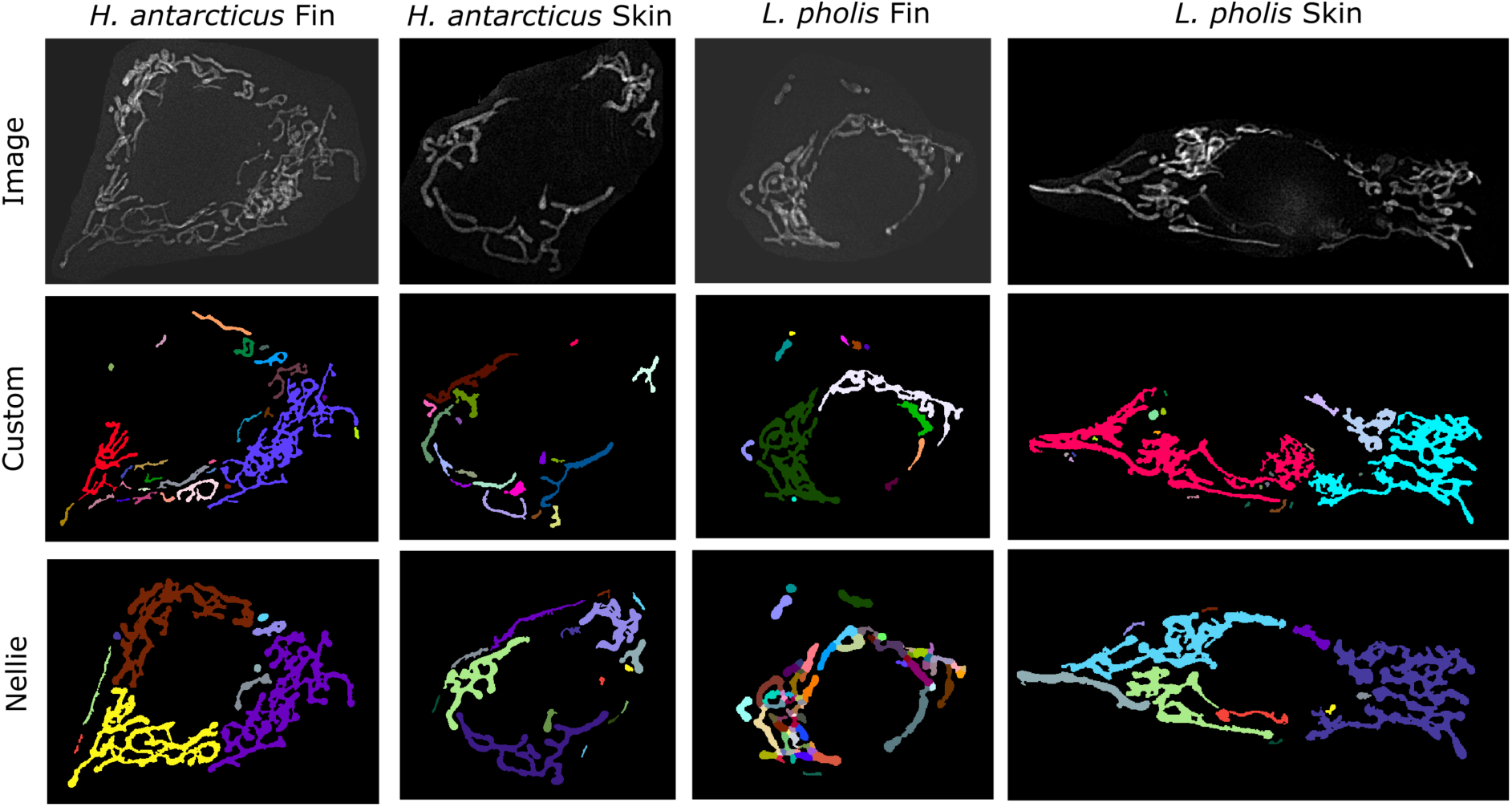
Nellie and a custom brightness-based segmentation method both perform accurate thresholding (minimising loss of mitochondrial area and inclusion of background), but imperfect segmentation of overlapping mitochondria.

**Supplementary Figure 7.**
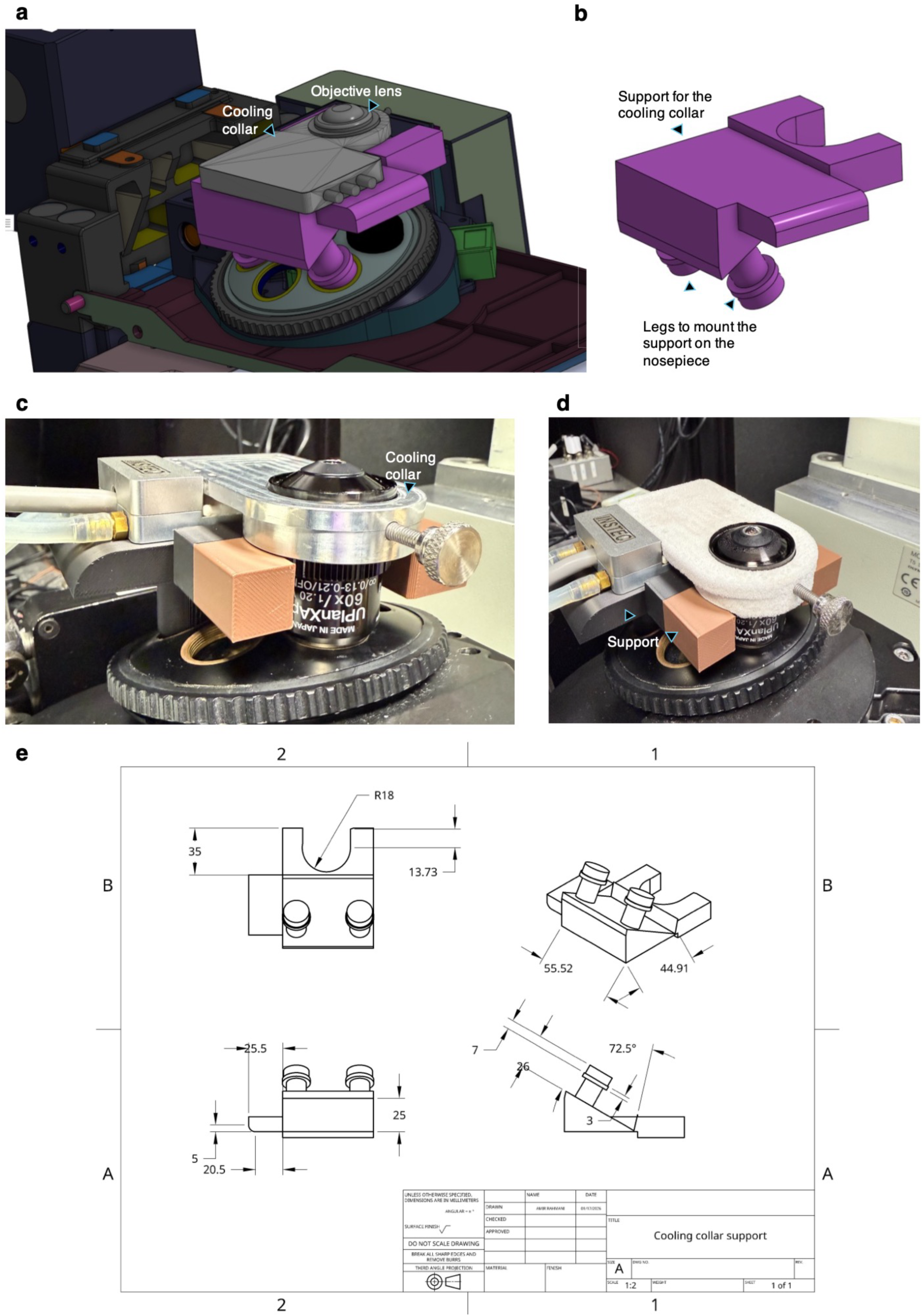
An enhanced cooling collar support further stabilises imaging. a) CAD rendering showing the integration of the cooling collar with the objective lens and the custom support mounted on an Olympus IX83 microscope nosepiece. b) CAD model of the cooling-collar support, highlighting the upper cradle that holds the collar and the lower legs used to mount the support onto the nosepiece. c-d) Photograph of the assembled cooling collar mounted on the objective lens with the custom support in place, providing increased mechanical stability during imaging c) at the start of imaging and d) after 3 h of operation at −13 °C, showing frost formation on the cooling collar. e) Technical drawing of the cooling-collar support with key dimensions and angles used for fabrication.

## 6 Methods

### 6.1 Animal collection and ethics statement

All *H. antarcticus* used in the experimental work were collected at Rothera Research Station, Adelaide Island, Antarctic Peninsula (67°34′07″S, 68°07′30″W) by SCUBA divers during the austral summer. Fish were collected under a permit (BAS-S7-2022/01) granted under Section 7 of the Antarctic Act 1994. The fish were returned to the UK and maintained in a recirculating flow aquarium in Cambridge, under a 12 h light/dark regime (water temperature 0.0±1.0 °C, salinity 34–36 PSU). The fish were fed twice weekly to satiation on white fish and shelled krill (*Euphausia superba*).

All *L. pholis* used in this experimental work were collected in the intertidal zone at low tide in the Menai Strait (Anglesey, Wales, UK) (53.23°N, 4.16°W) and transported to the experimental facility in Cambridge by vehicle, in cooled aquaria. The water temperature of the *L. pholis* stock tank was 12-15 °C, salinity was 34-36 PSU and a 12 h light/dark regime was imposed.

For both species, all adult fish were killed according to Home Office UK schedule 1 requirements before dissections were performed, at the British Antarctic Survey, Cambridge, UK.

### 6.2 Dissection of adult fish

Immediately after killing of adult fish, external tissues (fin and skin) were briefly swabbed with 70% ethanol. Dissection was then quickly performed on a bench, with the following tissues being collected: fin, skin, gonad, liver, muscle, kidneys, heart, and gills. Selected tissues were then used for cell culture or whole-tissue imaging. All tissues not used for any of these purposes were immediately flash-frozen by plunging into liquid nitrogen and were subsequently stored at -80 °C for use in non-cell culture experiments. Fish sex, length, and weight were recorded.

Following dissection, tissues to be used for cell culture were transferred immediately to ice-cold Leibovitz’s L-15 (Gibco, 21083-027) with antimicrobials and kept on ice. The antimicrobials were: PSN (Penicillin, Streptomycin, Neomycin) (Gibco, 15640-055) at standard concentration, Gentamicin (Sigma-Aldrich, G1272) at 0.05 mg/mL, and Amphotericin B (Sigma-Aldrich, A2942) at 2.5 μg/mL. Adult tissues trialled for *H. antarcticus* cell culture were fin, skin, ovary, liver, muscle, gill, heart and kidney. Embryo tissue was also trialled (see below, Fish cell culture). Only fin, skin, ovary and embryo were retained as suitable options for 2D cell culture.

### 6.3 Fish cell culture

Following dissection, tissue samples were transferred to a microbiological safety cabinet, where they were kept on ice throughout the culture process. Tissues were first washed in fresh L-15 with antimicrobials, with fin and skin being briefly scraped with a scalpel to remove external mucus. Tissues were then transferred to a Sylgard dish. Fin and skin were pinned down and cut into small fragments of approximately 2 mm by 2 mm to 3 mm by 3 mm using dissection scissors, with forceps being used to hold any unpinned tissue strips during cutting. Following cutting into small fragments, tissues were immediately transferred to 8-well dishes (Nunc™ Lab-Tek™ II Chambered Coverglass with No. 1.5 borosilicate glass bottom; Thermo Scientific, 155409) that had been pre-coated with 0.1% fish gelatin (Sigma-Aldrich, G7041) in MilliQ (autoclaved for sterilisation) for 30 min and had then been allowed to dry. For skin, explants were placed with the animal-internal side facing down (recognisable by its paler colour) and gently pressed down with forceps to ensure they were flat. Two pieces of tissue were placed per well for skin, and three per well for fin. All explants were left in closed dishes for 30 min to allow tissue adherence without drying out. 300 μL of culture medium was then added, which consisted of L-15 medium supplemented with 10% foetal bovine serum (FBS; (Sigma-Aldrich, F9665)) and the same antimicrobials as listed previously. For ovary, tissue was cut along its longest axis and opened, releasing the oocytes. The ovary membrane was then scraped and cut into small sections of approximately 2 mm by 2 mm to 3 mm by 3mm. These sections, along with 100 μL of any cells suspended in solution following scraping were collected and added to 0.1% fish gelatin-coated 8-well dishes. 200 μL of culture medium was then added.

Cell culture of embryos was performed in a similar manner as for adult tissues. Embryos were dechorionated on ice using watchmaker’s forceps in salt water from the aquarium. They were then transferred to L-15 with antimicrobials for washing, and subsequently to a Sylgard dish (with L-15 and antimicrobials) for dissection in a microbiological safety cabinet. Embryos were cut into 2 by 2 mm to 3 by 3 mm pieces using dissection scissors, with forceps being used to hold embryos to aid this process. These embryo fragments were then transferred to gelatin-coated 8-well LabTek dishes and left to adhere as previously, followed by addition of culture medium.

Tissue explants were left to grow for up to three weeks before imaging and were kept in an incubator at 2 °C (*H. antarcticus*) or 10 °C (*L. pholis*). Half of the culture medium was replaced with fresh medium twice per week, with all work being carried out on ice for *H. antarcticus*. For scoring culture success, explant outgrowth was evaluated using a brightfield microscope after one week; all explants that showed growth were considered to have been successful. As *L. pholis* cultures produced many loose clusters of cells, it was challenging to ascertain whether cells were derived from one of the tissue explants in a well, or from multiple. Culture success rates were therefore quantified per well rather than per explant for both species to ensure consistency. During fluorescence microscopy, only explants that showed good outgrowth free from obvious signs of cell stress such as large vacuoles or extensive cell rounding were included.

For optimisation of the culture, additional reagents were tested that were not included in the final protocol. For coating of culture dishes, these were poly-L-lysine (PLL) (Sigma-Aldrich, P4707), poly-D-lysine (PDL) (Gibco, 16021412), and PLL with laminin (Roche, 11243217001). PLL and PDL coating were performed for one hour, followed by washing. Laminin was then added to PLL-coated dishes and incubated overnight, followed again by washing. For the culture medium recipe, additional trialled reagents were salmon serum (Salmonics, 015), as a substitute for FBS, and NaCl (Thermo Scientific, J21618.A1), to increase osmolarity. For osmolarity adjustment trials, osmolarity was controlled by supplementing the cell culture medium with 8.276 mg/mL NaCl to make the osmolarity equivalent to that of sea water (600 mOsm/L). Final osmolarities were validated using a VAPRO 5600 vapor pressure osmometer. Media were then resterilised using a 0.2 μm membrane filter (Fisher Scientific, 15206869).

### 6.4 Mammalian cell culture

COS-7 cells were grown in T75 flasks (Greiner, 658175) or T25 flasks (Sarstedt, 83.3910.002) by incubation at 37 °C in a 5% CO_2_ atmosphere under controlled humidity. Complete medium for normal cell growth consisted of 90% high-glucose Dulbecco’s modified Eagle’s medium (DMEM) (Sigma-Aldrich, D6546), 10% FBS (Gibco, A5256801), 1% GlutaMAX (Gibco, 35050), and 1% Antibiotic-Antimycotic (Gibco, 15240). Splitting was performed at >80% confluency and medium was refreshed every 3–4 days. 24 hours ahead of experiments, 10K cells were seeded in each chamber of an 8-chamber glass well plate (Ibidi, 80807) with 200 µL of complete medium.

### 6.5 Sample preparation for fluorescence microscopy

#### 6.5.1 Fixed-tissue labelling and mounting

To obtain images of microridges *in situ* for *H. antarcticus* native tissue, freshly dissected fin and skin were used. Chunks of these tissues were immediately transferred to 1.5 mL Eppendorf tubes with L-15 on ice following dissection. Tissues were then briefly transferred to petri dishes with L-15 on ice, scraped to remove mucus, and cut into 3 mm by 3 mm squares. They were then returned to a 1.5 mL Eppendorf tube with L-15, on ice. The L-15 was then replaced with pre-chilled 4% methanol-free paraformaldehyde (PFA) (Thermo Scientific, 28906). Fixation was performed on ice for 30 min, after which samples were transferred to a rolling shaker at 4 °C for overnight fixation. Samples were subsequently washed three times with phosphate-buffered saline (PBS). To stain F-Actin, no further permeabilization step was performed; samples were incubated with Phalloidin-Atto647 (Sigma-Aldrich, 65906) overnight (approx. 50 nM final concentration), then washed again three times with PBS. Tissues were then immediately prepared for imaging by mounting in PBS on microscope slides (Marienfeld, 0704002) using Frame-Seal incubation chambers (Bio-Rad Laboratories) and 18×18 mm cover glass (Marienfeld, 0107032), with the external side of the tissue facing the coverglass for skin.

#### 6.5.2 Fixed-cell labelling

For imaging of F-actin in cells, fish cell cultures were washed once with PBS on ice (by replacing half the solution, avoiding the meniscus touching the sample), then immediately incubated in pre-chilled 2% PFA with 7.5% (w/v) sucrose in PBS for 30 min on ice (2% PFA final concentration achieved by removing half of culture medium and re-adding 4% PFA). This was followed by five washes with PBS. Samples were then permeabilised using 0.5% Triton-X (Sigma-Aldrich, T8787) at room temperature (RT) in PBS for 5 min, again followed by five washes with PBS (3 quick washes, then 2 times 5 min). Samples were then incubated with Phalloidin-Atto647, SYTO11, and Hoechst in PBS for 30 min at RT, followed by 5 washes in PBS (3 quick washes, then 2 times 5 min). Dye product information and concentrations used are summarised in Table 1.

**Table 1.**
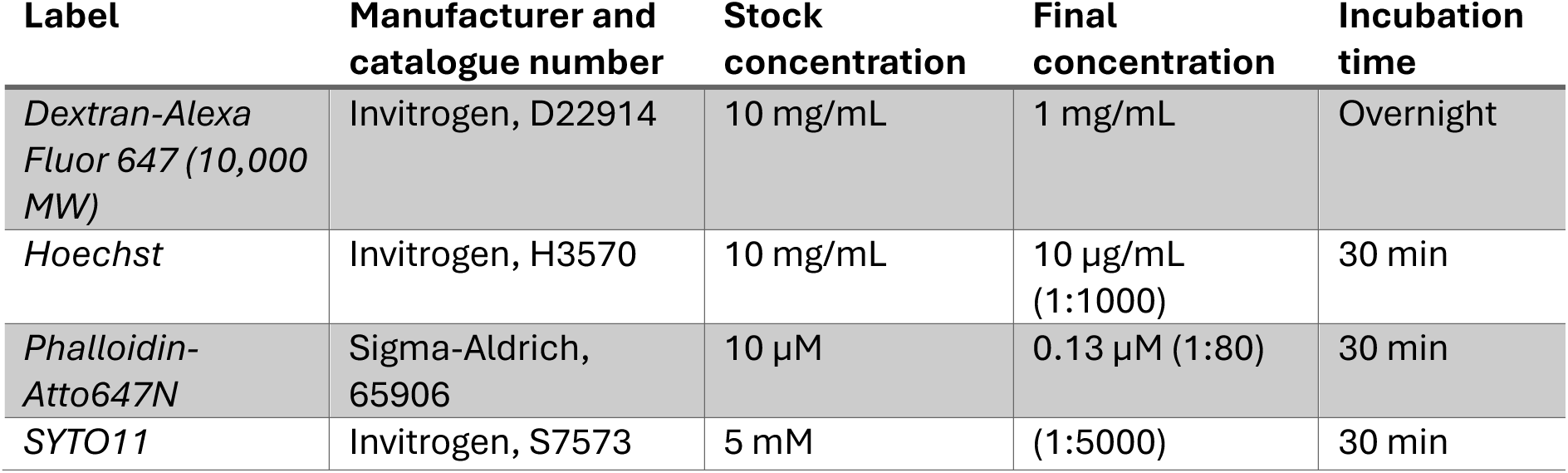
Labels for fixed-cell imaging.

For visualisation of Dextran internalisation, cultures were first washed twice with L-15. Dextran-Alexa647 in L-15 was then added to the cultures (see Table 1 for product information and concentration). After overnight incubation, cultures were washed seven times with L-15. They were then fixed as previously (without permeabilisation), stained with Hoechst to label nuclei, and imaged immediately.

#### 6.5.3 Live-cell labelling with dyes

For labelling of live cells, fish cells were washed once with L-15 (by replacing half the culture medium). Then, half the medium was again replaced with L-15 containing the fluorescent dye or dyes. After dye incubation, washing was performed by replacing half the medium in the well with fresh culture medium five times. Staining protocols are summarised in Table 2. For experiments involving multiple labels, dyes were incubated simultaneously. Dye incubations were all carried out on ice and in the dark.

**Table 2.** Labels for live-cell imaging.

| **Dye** | **Manufacturer and catalogue number** | **Stock concentration** | **Final concentration** | **Incubation time** |
| --- | --- | --- | --- | --- |
| *DRAQ7* | Biostatus, DR70250 | 0.3 mM | 0.9 µM (3:1000) | 15 min |
| *Hoechst* | Invitrogen, H3570 | 10 mg/mL | 10 µg/mL (1:1000) | 15 min |
| *LysoTracker Red DND-99* | Invitrogen, L7528 | 1 mM | 1 µM (1:1000) | 15 min |
| *Nile Red* | Invitrogen, N1142 | 1 mg/mL | 1 µg/mL (1:1000) | 1 h |
| *MitoTracker Deep Red FM* | Invitrogen, M22426 | 1 mM | 1 µM (1:1000) | 15 min |
| *MitoTracker Orange CMTMRos* | Invitrogen, M7510 | 1 mM | 0.5 μM (1:2000) | 15 min |
| *SYTO12* | Invitrogen, S7574 | 4 mM | 1 μM (1:4000) | 15 min |
| *SYTO24* | Invitrogen, S7559 | 5 mM | 1 μM (1:5000) | 15 min |

For staining of mammalian cells, all medium was removed and replaced with fresh medium containing the labels of interest. Incubation was carried out within an incubator at 37 °C, 5% CO_2_ and controlled humidity. Concentrations and incubation times were the same as in Table 2.

#### 6.5.4 Transfection

Skin cell culture samples were washed 5 times with L-15 without serum or antibiotics to remove RNAse from the culture dish. RNA lipoplexes were formed by diluting Lipofectamine MessengerMax (ThermoFisher, LMRNA001) in Opti-MEM (ThermoFisher, 31985062) to a final concentration of 3% (v/v) and separately diluting GFP mRNA (Scientific Laboratory Supplies, MRNA11-100) to a final concentration of 4.5% (v/v) (250% of the recommended dosage). Both dilutions were incubated at room temperature for 10 minutes and then mixed at a 1:1 ratio and left to complex at room temperature for a further 5 minutes. Complexes were then added to culture wells (in L-15 without antibiotics or serum) at 20% of the total well volume. Cells were incubated in the dark at 2 °C for 11 days before imaging.

### 6.6 Fluorescence microscopy

#### 6.6.1 Imaging at controlled temperature

For imaging live fish samples at sub-ambient temperatures, a custom cooling collar was used, as described in Marty, Ward, Lamb, van Tartwijk, Peck, Clark and Kaminski ^17^. In brief, a Peltier-based cooling system was attached to the objective lens and the temperature in the field of view was calibrated. To avoid condensation at the back aperture of the objective lens, it was periodically dried with nitrogen gas using a tube connected to a 3D-printed inlet. To improve mechanical stability and compatibility with the microscope body relative to the previous implementation ^17^, a rigid support was designed for the cooling collar. This support (Supplementary Figure 7) fits directly into the microscope nosepiece and enables secure coupling of the cooling collar to the objective lens without introducing lateral forces. This prevents any risk of the weight of the collar dragging or tilting the objective lens (which could lead to potential degradation of the lens with prolonged use).

Mammalian COS-7 cells were imaged in a microscope stage top micro-incubator (Okolab) with continuous air supply (37 °C and 5% CO_2_).

#### 6.6.2 Widefield microscopy

Widefield imaging was performed using a custom microscope. The microscope frame (IX83, Olympus) was equipped with a four-channel LED head (peak excitation wavelengths of 405 nm, 470 nm, 590 nm, and 625 nm; LED4D126, Thorlabs) and driver (DC4100, Thorlabs), a set of excitation/emission filters for each channel (Table 3), and a scientific complementary metal-oxide semiconductor (sCMOS) camera (Zyla 4.2, Andor). The system was controlled with the software Micro-Manager ^56^. Images were acquired with the following objective lenses: a 1.25×/0.04 numerical aperture (NA) air objective lens (PlanApo N, Olympus), a 10×/0.25 NA air objective lens (Plan N, Olympus), a 60×/1.42 NA oil objective lens (PlanApo U, Olympus), or a 100×/1.40 NA oil objective lens (UPlanSApo, Olympus).

**Table 3.** Filter sets used for widefield microscopy.

| Channel | Manufacturer and product number(s) | Excitation band (nm) | Emission band (nm) | Dichroic cut-on (nm) |
| --- | --- | --- | --- | --- |
| *DAPI* | Semrock, DAPI-1160B-OFX | 382 – 393 | 417 – 477 | 409 |
| *GFP* | Thorlabs, TLV-UFF-GFP | 469 ± 17.5 | 525 ± 19.5 | 597 |
| *YFP* | Edmund Optics, 86-833 | 513 – 556 | 570 – 613 | 562 |
| *mCherry* | Thorlabs, TLV-U-FF-MCHC | 562 ± 20 | 641 ± 37.5 | 593 |
| *Cy3* | Thorlabs, TLV-UFF-Cy3.5 | 565 ± 12 | 620 ± 26 | 590 |
| *Alexa647* | Semrock, FF01-609/54, FF01-698/70, FF660-Di02 | 582 – 636 | 662 – 732 | 660 |

#### 6.6.3 Structured illumination microscopy (SIM)

SIM was performed using a custom three-colour system built around an Olympus IX71 microscope body, as described in Young, Ströhl and Kaminski ^57^. Laser wavelengths of 488 nm (iBEAM-SMART-488, Toptica), 561 nm (OBIS 561, Coherent), and 640 nm (MLD 640, Cobolt) were used to excite the fluorophores in the samples. The laser beam was expanded to fill the display of a ferroelectric binary Spatial Light Modulator (SLM) (SXGA-3DM, Forth Dimension Displays) to pattern the light with a grating structure. The polarisation of the light was controlled with a Pockels cell (M350-80-01, Conoptics). A 60×/1.2 NA water immersion objective lens (UPLXAPO60XW, Evident) focused the structured illumination pattern onto the sample. This objective lens also collected the fluorescence emission from the sample, which, after passing through the tube lens and appropriate emission filters (FF01-525/30, FF01-600/37, FF01-676/29, all from Semrock), was projected onto an sCMOS camera (C11440, Hamamatsu). The maximum laser power density at the sample was 36.9 W/cm^2^ for the 488 nm laser, 52.0 W/cm^2^ for the 561 nm laser, and 26.4 W/cm^2^ for the 640 nm laser. Raw images were acquired with the HCImage software (Hamamatsu) to record image data to disk and a custom LabView program (freely available upon request) to synchronise the acquisition hardware. Multicolour images were registered by characterising channel displacement using a matrix generated with TetraSpeck beads (Life Technologies).

### 6.7 Image analysis

#### 6.7.1 Nile Red images

For each cell, a three-channel set of four images was taken at a 0-ms interval, ensuring negligible sample movement between frames (excitation at 488 nm, 561 nm, and 640 nm). A median-projection image was then made for each channel to reduce noise. Using the 600/37 channel (showing all organelles), individual cells were masked using the ‘Clear Outside’ function in Fiji ^58^. The channels were then separated into images for further analysis. For analysis of the 525/30 channel (lipid droplets), the following actions were performed: binarisation through manual thresholding, a watershed step to separate near-touching particles, and particle analysis using Fiji’s ‘analyse particles’ function (with a minimum particle size of 0.01 μm^2^ and the option to fill holes enabled). For each cell, this returned the number of droplets above the size threshold, their total area, and average area, as well as an area measurement for each individual droplet. For the 676/29 channel (perinuclear bodies), the same procedure was followed, but the minimum particle size was 0.1 μm^2^ and a watershed step was not included (due to the internal heterogeneity in staining of vesicles).

#### 6.7.2 Mitochondrial morphology

Mitochondria were segmented using Nellie ^34^, an automated deep learning-based image analysis tool implemented as a plugin within the napari image analysis platform ^59^. Images were restricted to single cells; where mitochondria from neighbouring cells were present in the field of view, those image regions were removed prior to segmentation using the ‘Clear Outside’ function in Fiji ^58^. The resulting images were then loaded into napari and analysed using the Nellie workflow. The model was applied to maximum intensity projected images. For each image, Nellie automatically detected individual mitochondrial objects and extracted morphological and intensity-based features.

To quantify the mitochondrial network at the cellular level, we developed a custom Python script that processed Nellie’s per-organelle output (features_organelles.csv) to calculate per-cell metrics. For each cell, we computed: (i) mitochondrial count (ii) total mitochondrial area; and (iii) form factor [(perimeter^2^)/(4π·surface area)]. However, due to imperfect segmentation, only the mitochondrial area measurement was used in subsequent analysis steps. The script processed all experimental images in batch mode, generating both individual and combined output files.

For mitochondrial segmentation and quantification using an automated thresholding method, we built a custom batch-processing macro implemented in Fiji/ImageJ. The same images masked to contain single cells were used as for analysis with Nellie. Firstly, the channel containing MitoTracker signal was isolated from any other channels. A background subtraction was then performed using a rolling-ball radius of 10 px (approximately 0.5 μm), reducing the background noise and artifacts from reconstructed SIM images. Automated thresholding was then applied using the Huang algorithm ^60^, which works by minimising the fuzziness of the given image. Compared to the commonly used Otsu thresholding, this method produces smoother and more stable thresholds in noisy or limited-contrast images. An optional manual thresholding step is added afterward, which can further optimise the segmentation performance if needed. After thresholding, images were computed into binary masks and further refined using morphological closing and dilation to improve the mitochondria continuity and reduce small segmentation holes. Each individual mitochondrion mask that was less than 50 px^2^ (approximately 0.14 μm^2^) was removed and the rest of the masks were analysed using the “Analyze Particles” function in Fiji, to exclude noise and artifacts. For each image, mitochondrial morphological parameters statistics including mitochondrion numbers, average mitochondrion area, perimeter, and form factor were automatically exported for further analysis. Total mitochondrial area per cell was the only parameter used in subsequent comparison with Nellie’s output.

#### 6.7.3 Mitochondrial dynamics

Single-channel timelapses of mitochondria in live cells (acquired at 3-s intervals) were pre-processed using the ‘Enhance Contrast > Normalize > Process all n slices > use stack histogram’ function in Fiji. Normalised stacks were then uploaded into the Mitometer application in MATLAB. Custom ‘Advanced settings’ were used, namely a Custom gauss filter sigma set to 15 and a Custom threshold set to 10. Processed images were individually assessed for quality. All the tracks not corresponding to a mitochondrion were removed and only tracks confidently assessed as accurate with a track length of 2 or higher were retained and exported for analysis.

### 6.8 Statistical analyses

Each comparative analysis between species and/or tissue types included at least two biological replicates. A single biological replicate included all data derived from the same individual fish, from multiple technical replicates (distinct explant cultures). Specific tests used for each analysis are given below with relevant justification.

For all analyses of images of Nile Red-stained cells (both 525/30 and 676/29 emission channels) and all comparisons in Figure 2, statistical comparisons between *H. antarcticus* and *L. pholis* were performed using Mann-Whitney test, with significance set at *p*<0.05. Effect sizes were reported as Cohen’s *d*. Normality was assessed using the Shapiro-Wilk test and variances were evaluated using Levene’s test. All data were non-normally distributed except for *H. antarcticus* perinuclear body area and variance was unequal for all comparisons between *H. antarcticus* and *L. pholis*.

For mitochondrial morphology and mitochondrial dynamic analyses, all statistical comparisons between species and between tissue types were performed using Mann-Whitney U test, with significance set at *p*<0.05. Non-parametric testing was selected following assessment of normality and equality of variances. Effect sizes were reported as Cohen’s *d*.

All statistical analyses were performed in Python (SciPy library).

## References

1. Knapp, B.D. & Huang, K.C. The effects of temperature on cellular physiology. Annual Review of Biophysics 51, 499–526 (2022).

2. van Tartwijk, F.W. & Kaminski, C.F. Protein condensation, cellular organization, and spatiotemporal regulation of cytoplasmic properties. Advanced Biology 6, 2101328 (2022).

3. Clarke, A. & Crame, J.A. The Southern Ocean benthic fauna and climate change: a historical perspective. Philosophical Transactions of the Royal Society B: Biological Sciences 338, 299–309 (1992).

4. Peck, L.S., Convey, P. & Barnes, D.K. Environmental constraints on life histories in Antarctic ecosystems: tempos, timings and predictability. Biol Rev Camb Philos Soc 81, 75–109 (2006).

5. Peck, L.S. A cold limit to adaptation in the sea. Trends in Ecology & Evolution 31, 13–26 (2016).

6. Peck, L.S. Antarctic marine biodiversity: adaptations, environments and responses to change. Oceanography and Marine Biology (2018).

7. Fraser, K.P., Clarke, A. & Peck, L.S. Low-temperature protein metabolism: seasonal changes in protein synthesis and RNA dynamics in the Antarctic limpet Nacella concinna Strebel 1908. J Exp Biol 205, 3077–3086 (2002).

8. Fraser, K.P.P., Clarke, A. & Peck, L.S. Growth in the slow lane: protein metabolism in the Antarctic limpet Nacella concinna (Strebel 1908). Journal of Experimental Biology 210, 2691–2699 (2007).

9. Fraser, K.P.P., Peck, L.S., Clark, M.S. & Clarke, A. A comparative study of tissue protein synthesis rates in an Antarctic, Harpagifer antarcticus and a temperate, Lipophrys pholis teleost. Comparative Biochemistry and Physiology Part A: Molecular & Integrative Physiology 295, 111650 (2024).

10. Fraser, K.P.P., Peck, L.S., Clark, M.S., Clarke, A. & Hill, S.L. Life in the freezer: protein metabolism in Antarctic fish. Royal Society Open Science 9, 211272 (2022).

11. Todgham, A.E., Hoaglund, E.A. & Hofmann, G.E. Is cold the new hot? Elevated ubiquitin-conjugated protein levels in tissues of Antarctic fish as evidence for cold-denaturation of proteins in vivo. Journal of Comparative Physiology B 177, 857–866 (2007).

12. Shin, S.C. et al. Transcriptomics and comparative analysis of three antarctic notothenioid fishes. PLoS One 7, e43762 (2012).

13. Hofmann, G.E., Lund, S.G., Place, S.P. & Whitmer, A.C. Some like it hot, some like it cold: the heat shock response is found in New Zealand but not Antarctic notothenioid fishes. Journal of Experimental Marine Biology and Ecology 316, 79–89 (2005).

14. Place, S.P. & Hofmann, G.E. Constitutive expression of a stress-inducible heat shock protein gene, hsp70, in phylogenetically distant Antarctic fish. Polar Biology 28, 261–267 (2005).

15. Clark, M.S., Fraser, K.P.P., Burns, G. & Peck, L.S. The HSP70 heat shock response in the Antarctic fish Harpagifer antarcticus. Polar Biology 31, 171–180 (2008).

16. Berthelot, C. et al. Adaptation of proteins to the cold in Antarctic fish: a role for methionine? Genome Biology and Evolution 11, 220–231 (2018).

17. Marty, A.-P.M. et al. A high-resolution microscopy system for biological studies of cold-adapted species under physiological conditions. Small Methods 9, 2401682 (2025).

18. Peck, L.S., Clark, M.S., Morley, S.A., Massey, A. & Rossetti, H. Animal temperature limits and ecological relevance: effects of size, activity and rates of change. Functional Ecology 23, 248–256 (2009).

19. Clark, M.S. et al. Lack of long-term acclimation in Antarctic encrusting species suggests vulnerability to warming. Nature Communications 10, 3383 (2019).

20. LeClair, E.E. The last half century of fish explant and organ culture. Zebrafish 18, 1–19 (2021).

21. Bilyk, K.T. & DeVries, A.L. Heat tolerance and its plasticity in Antarctic fishes. Comparative Biochemistry and Physiology Part A: Molecular & Integrative Physiology 158, 382–390 (2011).

22. Stanwell-Smith, D. & Peck, L.S. Temperature and embryonic development in relation to spawning and field occurrence of larvae of three Antarctic echinoderms. The Biological Bulletin 194, 44–52 (1998).

23. Depasquale, J.A. Actin microridges. The Anatomical Record 301, 2037–2050 (2018).

24. Campbell, K. & Casanova, J. A common framework for EMT and collective cell migration. Development 143, 4291–4300 (2016).

25. Greenspan, P., Mayer, E.P. & Fowler, S.D. Nile red: a selective fluorescent stain for intracellular lipid droplets. J Cell Biol 100, 965–973 (1985).

26. Dutta, A.K., Kamada, K. & Ohta, K. Spectroscopic studies of nile red in organic solvents and polymers. Journal of Photochemistry and Photobiology A: Chemistry 93, 57–64 (1996).

27. Zhanghao, K. et al. High-dimensional super-resolution imaging reveals heterogeneity and dynamics of subcellular lipid membranes. Nature Communications 11, 5890 (2020).

28. Liu, H., Tang, H. & Pang, S. Role of lysosomal morphology in aging and age-related diseases. Ageing Research Reviews 115, 103033 (2026).

29. Chazotte, B. Labeling lysosomes in live cells with LysoTracker. Cold Spring Harbor Protocols 2011, pdb. prot5571 (2011).

30. Katz, M.L. & Robison, W.G. What is lipofuscin? Defining characteristics and differentiation from other autofluorescent lysosomal storage bodies. Archives of Gerontology and Geriatrics 34, 169–184 (2002).

31. Klumperman, J. & Raposo, G. The complex ultrastructure of the endolysosomal system. Cold Spring Harb Perspect Biol 6, a016857 (2014).

32. Guderley, H.E. & Pierre, J.S. in Animals and Temperature: Phenotypic and Evolutionary Adaptation. (eds. I.A. Johnston & A.F. Bennett) 127–152 (Cambridge University Press, Cambridge; 1996).

33. Neikirk, K. et al. MitoTracker: a useful tool in need of better alternatives. European Journal of Cell Biology 102, 151371 (2023).

34. Lefebvre, A.E.Y.T. et al. Nellie: automated organelle segmentation, tracking and hierarchical feature extraction in 2D/3D live-cell microscopy. Nature Methods 22, 751–763 (2025).

35. Walesby, N.J. & Johnston, I.A. Fibre types in the locomotory muscles of an Antarctic teleost, Notothenia rossii: A histochemical ultrastructural and biochemical study. Cell and tissue research 208, 143–164 (1980).

36. Johnston, I.A., Calvo, J., Guderley, H., Fernandez, D. & Palmer, L. Latitudinal variation in the abundance and oxidative capacities of muscle mitochondria in perciform fishes. Journal of Experimental Biology 201, 1–12 (1998).

37. Lefebvre, A., Ma, D., Kessenbrock, K., Lawson, D.A. & Digman, M.A. Automated segmentation and tracking of mitochondria in live-cell time-lapse images. Nat Methods 18, 1091–1102 (2021).

38. Begeman, A., Smolka, J.A., Shami, A., Waingankar, T.P. & Lewis, S.C. Spatial analysis of mitochondrial gene expression reveals dynamic translation hubs and remodeling in stress. Science Advances 11, eads6830 (2025).

39. Stephan, T., Roesch, A., Riedel, D. & Jakobs, S. Live-cell STED nanoscopy of mitochondrial cristae. Scientific Reports 9, 12419 (2019).

40. Ren, W. et al. Visualization of cristae and mtDNA interactions via STED nanoscopy using a low saturation power probe. Light: Science & Applications 13, 116 (2024).

41. Houghton, O.H., Mizielinska, S. & Gomez-Suaga, P. The interplay between autophagy and RNA homeostasis: implications for amyotrophic lateral sclerosis and frontotemporal dementia. Frontiers in Cell and Developmental Biology Volume 10 - 2022 (2022).

42. Wang, D.W., Peng, Z.J., Ren, G.F. & Wang, G.X. The different roles of selective autophagic protein degradation in mammalian cells. Oncotarget 6, 37098–37116 (2015).

43. Tondera, D. et al. SLP-2 is required for stress-induced mitochondrial hyperfusion. EMBO Journal 28, 1589–1600 (2009).

44. Liu, Y.J., McIntyre, R.L., Janssens, G.E. & Houtkooper, R.H. Mitochondrial fission and fusion: a dynamic role in aging and potential target for age-related disease. Mechanisms of Ageing and Development 186, 111212 (2020).

45. Ježek, J., Cooper, K.F. & Strich, R. Reactive oxygen species and mitochondrial dynamics: the Yin and Yang of mitochondrial dysfunction and cancer progression. Antioxidants (Basel) 7 (2018).

46. Xu, X., Pang, Y. & Fan, X. Mitochondria in oxidative stress, inflammation and aging: from mechanisms to therapeutic advances. Signal Transduction and Targeted Therapy 10, 190 (2025).

47. Gomes, L.C., Benedetto, G.D. & Scorrano, L. During autophagy mitochondria elongate, are spared from degradation and sustain cell viability. Nature Cell Biology 13, 589–598 (2011).

48. Rambold, A.S., Kostelecky, B., Elia, N. & Lippincott-Schwartz, J. Tubular network formation protects mitochondria from autophagosomal degradation during nutrient starvation. Proceedings of the National Academy of Sciences 108, 10190–10195 (2011).

49. Sidell, B.D. Intracellular oxygen diffusion: the roles of myoglobin and lipid at cold body temperature. Journal of Experimental Biology 201, 1119–1128 (1998).

50. Czarnoleski, M. & Verberk, W.C.E.P. Cell size matters: a unifying theory across the tree of life. Trends in Ecology & Evolution (2025).

51. Verberk, W. et al. Shrinking body sizes in response to warming: explanations for the temperature-size rule with special emphasis on the role of oxygen. Biol Rev Camb Philos Soc 96, 247–268 (2021).

52. Pichon, X. et al. Visualization of single endogenous polysomes reveals the dynamics of translation in live human cells. The Journal of Cell Biology (2016).

53. Ströhl, F. et al. Single molecule translation imaging visualizes the dynamics of local β-actin synthesis in retinal axons. Scientific Reports 7, 709 (2017).

54. Lu, M. et al. ERnet: a tool for the semantic segmentation and quantitative analysis of endoplasmic reticulum topology. Nature Methods 20, 569–579 (2023).

55. Portner, H.-O., Bock, C. & Mark, F.C. Oxygen- and capacity-limited thermal tolerance: bridging ecology and physiology. Journal of Experimental Biology 220, 2685–2696 (2017).

56. Edelstein, A., Amodaj, N., Hoover, K., Vale, R. & Stuurman, N. Computer control of microscopes using μManager. Curr Protoc Mol Biol Chapter 14, Unit14.20 (2010).

57. Young, L.J., Ströhl, F. & Kaminski, C.F. A guide to structured illuminaOon TIRF microscopy at high speed with mulOple colors. JoVE, e53988 (2016).

58. Schindelin, J. et al. Fiji: an open-source plasorm for biological-image analysis. Nature Methods 9, 676–682 (2012).

59. Selzer, G.J. et al. napari-imagej: ImageJ ecosystem access from napari. Nature Methods 20, 1443–1444 (2023).

60. Huang, L.-K. & Wang, M.-J.J. Image thresholding by minimizing the measures of fuzziness. Pattern recognition 28, 41–51 (1995).

